# Hybrid-DIA: Intelligent Data Acquisition for Simultaneous Targeted and Discovery Phosphoproteomics in Single Spheroids

**DOI:** 10.1101/2022.12.19.520999

**Authors:** Ana Martínez-Val, Kyle Fort, Claire Koenig, Leander Van der Hoeven, Giulia Franciosa, Thomas Moehring, Yasushi Ishihama, Yu-ju Chen, Alexander Makarov, Yue Xuan, Jesper V. Olsen

## Abstract

Achieving sufficient coverage of regulatory phosphorylation sites by mass spectrometry (MS)-based phosphoproteomics for signaling pathway reconstitution is challenging when analyzing tiny sample amounts. We present a novel hybrid data-independent acquisition (DIA) strategy (hybrid-DIA) that combines targeted and discovery proteomics through an Application Programming Interface (API) to dynamically intercalate DIA scans with accurate triggering of multiplexed tandem MS scans of predefined (phospho)peptide targets. By spiking-in heavy stable isotope labeled phosphopeptide standards covering seven major signaling pathways, we benchmarked hybrid-DIA against state-of-the-art targeted MS methods (i.e. SureQuant) using EGF-stimulated HeLa cells and found the quantitative accuracy and sensitivity to be comparable while hybrid-DIA also profiled the global phosphoproteome. To demonstrate the robustness, sensitivity and potential of hybrid-DIA, we profiled chemotherapeutic agents in single colon carcinoma multicellular spheroids and evaluated the difference of cancer cells in 2D vs 3D culture. Altogether, we showed that hybrid-DIA is the way-to-go method in highly sensitive phospho-proteomics experiments.

## Introduction

Liquid chromatography tandem mass spectrometry (LC-MS/MS) acquisition strategies for proteomics can be divided into two main categories: discovery and targeted proteomics methods. The aim of discovery approaches is to achieve the most comprehensive coverage of the proteome or sub-proteome under investigation. Although still far from completeness, the most representative MS acquisition method to achieve this in single-shot analysis is data independent-acquisition (DIA)^1^. DIA has proven capable of maximizing the number of identifications obtained per sample, especially when studying post-translational modification (PTM) landscapes^2–4^. For example, DIA-based discovery proteomics is a powerful technology for studying global changes in the phosphoproteome^5,6^ in cells, tissues and organisms. Phosphoproteomics typically requires enrichment of phosphopeptides prior to MS analysis, limiting the scope of the analysis when sample availability is scarce. Fortunately, the phosphoproteomics technology has advanced significantly in recent years in terms of sensitivity and robustness as optimal amounts required for phosphopeptide-enrichment prior to MS analysis has been reduced by a factor of ten from ~2 mg to 200 ug of peptide input^7–9^. Yet, despite the latest boost in depth achieved by single-shot phosphoproteomics due to DIA, many biologically important phosphopeptide targets of low abundance are often missed in phosphoproteomics experiments. This shortcoming makes certain biological systems inaccessible to traditional phosphoproteomics analysis due to their inherent limited material, such as analysis of phosphoproteomes from single spheroids or organoids, tumor fine needle aspiration biopsies or even single-cells. In these scenarios, the protein amount available for phosphoproteomics analysis is sub-optimal, and in the best case scenario only allows for single-injection LC-MS/MS analysis. Therefore, for restricted biological matrices it is essential to enhance sensitivity of the analysis to maximize the phosphoproteome coverage in each MS run.

In distinction to discovery proteomics, targeted proteomics approaches provide improved detection and quantification of a predefined set of peptides with good accuracy and precision across multiple runs; however, single and parallel reaction monitoring (SRM/PRM) methods required extensive method optimization which, among others factors, limits the number of target peptides that can be accurately monitored. To address this limitation, intelligent acquisition methods have been developed, such as spike-in triggered PRM acquisition methods (i.e. SureQuant^10^, TOMAHAQ^11^, Pseudo-PRM^12^, in which targeted scans are triggered by detection of synthetic heavy labeled peptides spiked into the samples before MS analysis. Consequently, intelligent MS data acquisition methods able to combine discovery proteomics with targeted acquisition of selected peptides of interest would improve the sensitivity and reproducibility of phosphoproteomics, especially by ensuring accurate quantification of key phosphorylation pathway markers complementing the analysis of the global phosphoproteome footprint.

Translational scientists face a dilemma when having to choose between comprehensive discovery proteomics-based profiling and sensitive targeted quantitation, especially when analyzing large sample cohorts. Discovery proteomics is commonly used for biomarker identification, having a great potential for unveiling prognostic and predictive biomarkers; however, it still lacks the sensitivity to accurately quantify all the biomarkers of interest. Therefore, in the validation phase, targeted MS quantitation of the potential markers usually have to be employed. This leads to high cost, time loss and additional sample consumption. To address these challenges, we developed an intelligent MS data acquisition strategy termed hybrid-DIA that combines comprehensive proteome profiling via data-independent-acquisition mass spectrometry with simultaneous on-the-fly triggering of parallel reaction monitoring (PRM) and multiplexed MS/MS (MSx) scans for sensitive and accurate quantification of the predefined marker peptides. This hybrid DIA-MS acquisition strategy substantially increases throughput and coverage while reducing sample consumption. It presents a new capability to combine data-driven and hypothesis-driven MS acquisition approaches in one go. The hybrid-DIA acquisition strategy uses an Application Programming Interface (API) to dynamically intercalate DIA scans with multiplexed tandem MS scans of predefined (phospho)peptide targets by spiking-in heavy stable isotope labeled phosphopeptide standards. In this work, we benchmarked hybrid-DIA to show its benefits when compared to conventional DIA and triggered targeted proteomics acquisition methods. Furthermore, we demonstrated its potential to maximize the information retrievable from challenging low-level phospho-proteomics samples by using hybrid-DIA to describe the mechanism of action of the chemotherapeutic drug 5-fluorouracil (5-FU) in single colon cancer multicellular spheroids compared to conventional monolayer cell culture.

## Results

### Implementation of an API to enable targeted and discovery proteomics

Hybrid-DIA is an intelligent MS data acquisition strategy implemented through an Application Programming Interface (API) in the Tune software controlling an Orbitrap^™^ Exploris^™^ 480 mass spectrometer^13^ (Supplementary Data 1). The acquisition method combines a standard DIA acquisition scheme consisting of a full scan MS followed by a flexible number of MS/MS precursor isolation windows with on the fly triggering of intercalated multiplexed MS/MS (MSx) scans. Multiplex scans consist of spiked-in heavy stable isotopically labeled peptide standards (IS) and their predicted endogenous (ENDO) counterparts based on a predefined precursor inclusion list (Fig. 1A). Detection of IS peptide precursors in a full-scan MS triggers a fast low-resolution PRM-MS/MS scan of the observed IS peptides. Automatic matching of a minimum number of predefined fragment ions with high mass accuracy in the MS/MS of the heavy isotope labeled standards triggers additional multiplexed MS/MS spectra of each of the individual heavy IS co-analyzed with their corresponding endogenous peptide, respectively. The triggered multiplexed MS/MS scans of the co-isolated IS and ENDO peptides (IS/ENDO-MSx) are acquired with narrow quadrupole isolation window, high-resolution Orbitrap detection and differential ion injection times (IT) to equalize the precursor abundances (Fig. 1B). Thus, the detection and accuracy of quantitation for the endogenous peptide are maximized. All of the triggered targeted PRM and MSx scans are performed while simultaneously acquiring the conventional DIA data. As a result, hybrid-DIA raw files contain both unbiasedly acquired DIA data as well as a selection of targeted scans of peptides with higher sensitivity and better quantitative accuracy and precision. To process hybrid-DIA raw files, it is necessary to separate the DIA scans from the IS/ENDO-MSx scans. For retrieving the DIA data, we employed the HTRMS convertor tool co-installed with Spectronaut software, which was used to analyzed the DIA data. Moreover, we have developed an in-house analysis pipeline to extract the IS/ENDO-MSx scans and quantify all IS/ENDO peptide pairs detected. To do this, we first extract all IS/ENDO-MSx scans into a separate mzML^14^ file, whilst simultaneously extracting the information on the differential IT used in the MSx scans. Next, we load the mzML file into Skyline^15^ to readout the raw fragment ion intensities from each pair of IS/ENDO peptides. Finally, the resulting files are loaded into an R-shiny app that we have designed to correct for the differential injection time (Supplementary Fig. 2A), and determine the area-under-the-curve (AUC) of both ENDO and IS peptides. Additionally, the R-shiny app enables visual inspection of the MSx scans and the resulting quantification, scaled by conditions or as ENDO/IS ratios (Supplementary Fig. 2A-B, Supplementary Data 2).

**Figure 1.**
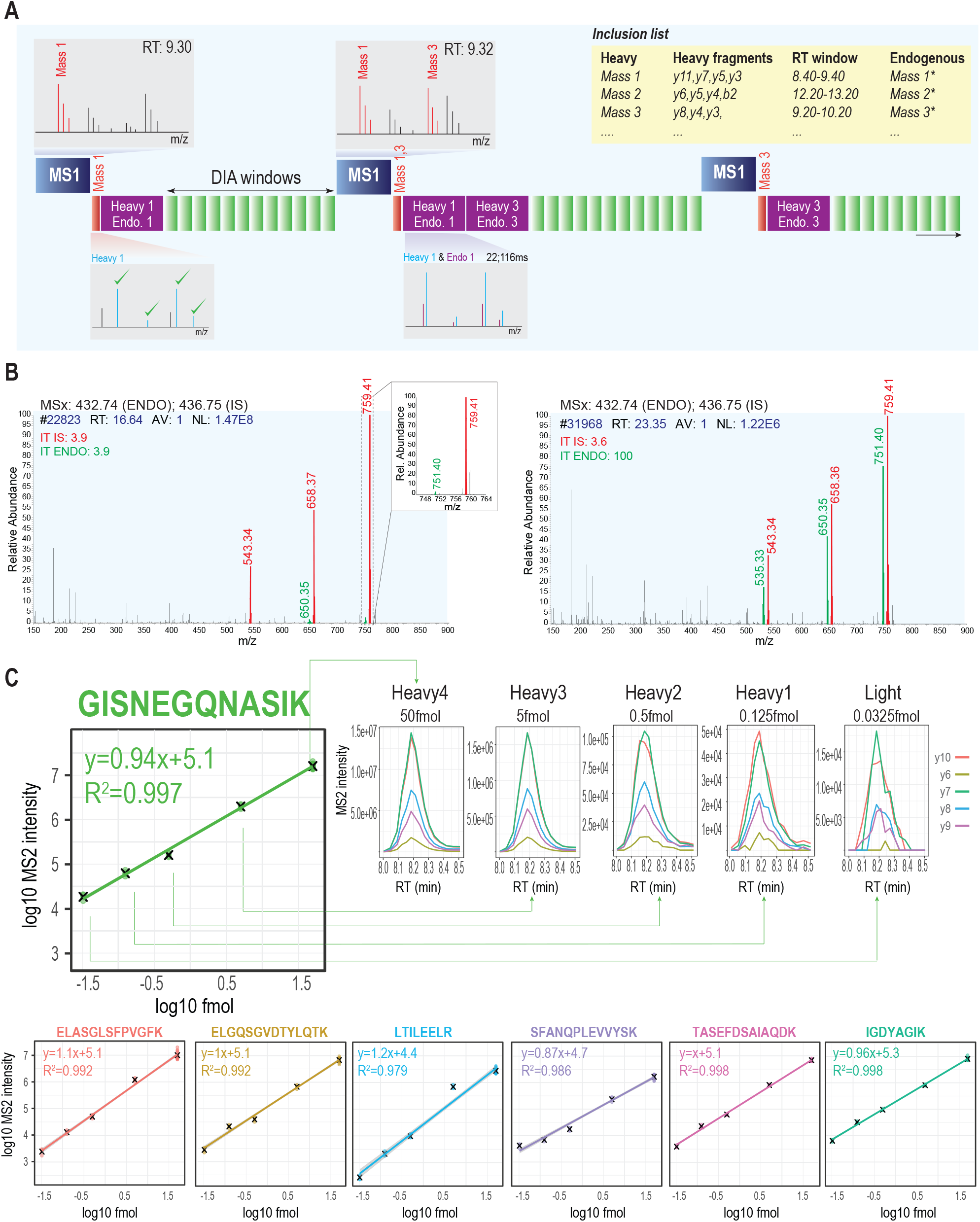
(A) Description of the novel hybrid-DIA acquisition method. (B) Example of the increase in sensitivity by using differential injection time in the multiplexed (MSx) scans. In red, spiked-in heavy stable isotopically labeled peptide standard (IS) and in green the triggered endogenous peptide (ENDO). (C) Demonstration on the accuracy, precision and linearity of the quantification in the MSx scans using the Pierce Suitability Standard Mix, which is comprised of 7 peptides, each one with 5 isotopologue sequences present in a dilution series ranging from 0.5pmol/ul to 0.3 fmol/ul. The isotopologue with the highest concentration (Heavy4) was used as a triggering peptide, and the subsequent peptides were used as “endogenous” counterparts, and were triggered sequentially in the acquisition cycle.

To initially validate the accuracy and sensitivity of the quantification results from the MSx scans derived from the hybrid-DIA files, we used the Pierce Suitability Standard mixture comprising 7 different peptides, each one with 5 isotopologue sequences^16^ present in a dilution series ranging from 0.5pmol/μl to 0.3 fmol/μl. We injected 0.1ul of the Pierce Suitability Standard mixture, and used the more abundant isotopologue (50 fmol on column) to trigger MSx scans of the remaining 4 isoforms (5, 0.5, 0.125 and 0.0325 fmol, respectively). Samples were analyzed using the Whisper flow technology 40 samples per day (SPD) gradient on an Evosep One LC system coupled to an Orbitrap Exploris 480 mass spectrometer operating with the hybrid-DIA API. Specifically, we acquired MSx scans using a maximum of 116 ms injection time and automatic gain control (AGC) target of 1e6. As a result, all seven peptides were correctly detected and all four isotopologues quantified for each peptide. Using our hybrid-DIA analysis pipeline (Supplementary Data 2) we quantified the intensities from each one of the isotopologues measured in the hybrid-DIA scan. We found that the targeted MSx scans allow to correctly quantifying amounts as low as 0.0325 fmol, and the quantification shows perfect linearity for the entire dynamic range covered (Fig. 1C).

### Hybrid-DIA improves the limit of detection and quantification of predefined targets in phosphoproteomics whilst preserving the coverage of the phosphoproteome

The spectral library-free directDIA MS analysis strategy has recently emerged as a high-throughput and straightforward approach for discovery-based phosphoproteomics^5^. However, such single-shot phosphoproteomes are still limited in coverage and are far from completeness^17^, and many phosphopeptides of interest might not be detected and quantified properly. Moreover, site-specific phosphorylation is a dynamic sub-stoichiometric post-translational modification (PTM) requiring specific phosphopeptide enrichment prior to MS analysis, which makes sample amounts available critical for effective phosphoproteomics analysis. Importantly, many sample types of biomedical interest are limited in protein amount (e.g. FACS sorted cells, FFPE or spheroids) restricting the possibility to perform both discovery proteomics and targeted MS validation from the same material. The hybrid-DIA methodology could alleviate the dilemma of choosing between DIA or PRM analysis, and thereby maximize the knowledge derived from a single sample, which is of special relevance for high-sensitivity phosphoproteomics applications.

To demonstrate the benefits of hybrid-DIA in terms of improved sensitivity, we benchmarked it against conventional DIA analysis in a cell line model for sensitive phosphoproteome analysis. Using A549 human lung adenocarcinoma cells, we performed phospho-enrichment from decreasing amounts of tryptic peptide digests, starting from 30 μg and down to 2.5 μg of peptide input. Samples were prepared in quadruplicates, each phosphopeptide-enrichment performed independently. To assess the potential of hybrid-DIA for measuring a predefined panel of phosphopeptides, we used the commercial SureQuant™ Multipathway Phosphopeptide Standard mixture containing 131 heavy-stable isotope labeled phosphopeptides of relevance covering seven major cellular signaling pathways. 50 fmol of the mixture was added to all samples, and subsequently half of them were analyzed in DIA mode, and half of them using the hybrid-DIA approach (Fig. 2A). The phosphopeptides contained in the SureQuant™ Multipathway Phosphopeptide Standard mixture are evenly distributed across the 20SPD chromatographic gradient (Supplementary Fig. 2A), and their endogenous counterparts are very diverse in MS signal intensities spanning several orders of magnitude. To prove the improved limit of detection of hybrid-DIA in the MSx scans, we extracted the ion chromatograms (XICs) of three peptides from the panel with difference abundances: AKT1S1:T246 (high abundance), TSC2:S939 (medium abundance) and PLCG1:Y783 (low abundance). Whilst the high abundance phosphopeptide (AKT1S1:T246) is clearly detected both in the MSx scans in hybrid-DIA mode and in the MSMS scans in standard DIA mode, it is clear that with reduced abundance the retrieved signal for both TSC2:S939 and PLCG1:Y783 is non-existent in standard DIA, but readily detected in MSx scans in hybrid-DIA, even at input amounts as low as 2.5 μg prior to phospho-enrichment (Fig. 2B).

**Figure 2.**
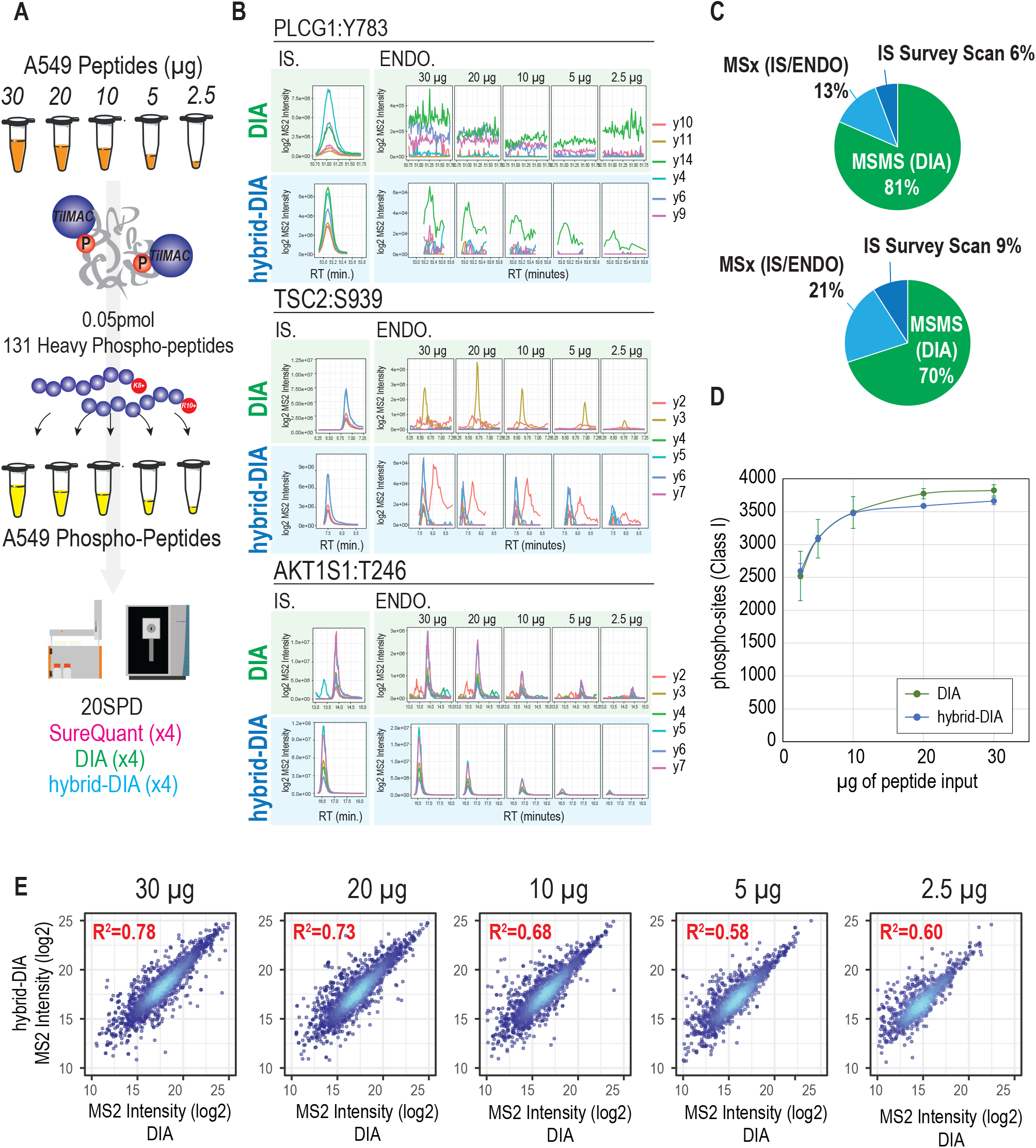
(A) Experimental design for benchmark analysis of hybrid-DIA versus standard DIA and SureQuant. Decreasing input material for phosphopeptides from A549 were used for phospho-enrichment, performed in quadruplicates per experimental condition (n=20, 4 replicates x 5 input amounts). Heavy labeled peptide mixture was added afterwards, and samples were analyzed either by hybrid-DIA, DIA or SureQuant. (B) XIC for three phosphopeptides spanning the dynamic range of the phosphopeptide mixture. In green the data from DIA runs, in blue the data from MSx scans in hybrid-DIA runs. MS2 intensity of hybrid-DIA runs has been corrected by injection time. (C) Pie charts representing the proportion of cycle time used by the hybrid-DIA API measured by number of scans (top) or in acquisition time (bottom). (D) Number of phospho-sites (class I) identified using Spectronaut (V17) in DIA (green, n=4) and hybrid-DIA (blue, n=4). (E) Correlation plot of quantified phospho-sites in hybrid-DIA runs (y-axis) versus DIA runs (x-axis) for the different dilutions. Correlation is indicated as R-squared.

One important concern that might arise when comparing standard DIA against hybrid-DIA is whether the inclusion of targeted scans during a normal DIA run will affect the cycle time significantly, and if so, impact the overall identification and quantification of peptides. To assess this, we calculated the percentage of measurement time employed by hybrid-DIA PRM and MSx scans in this experiment and found that almost 20% of the MS/MS scans are triggered by the API, comprising one third of the total MS/MS acquisition time not considering the full scans (Fig. 2C). However, most importantly, when comparing the number of phosphopeptides identified by directDIA using Spectronaut (v17) in each method, we did not see any notable difference between conventional DIA and hybrid-DIA runs (Fig. 2D). Furthermore, to evaluate the relationship between the total number of IS/ENDO targets on the inclusion list and the effect of DIA performance, we carried on an experiment targeting an increasing number of targeted phosphopeptides (50, 75 and 100 targeted peptides) (Supplementary Fig. 2A) with hybrid-DIA but decreasing the gradient to 40SPD (Supplementary Fig. 2B-C). As expected, an increase in the fraction of cycle time devoted to hybrid-DIA scans is observed, the more targets added to the inclusion (Supplementary Fig. 2B). This is accompanied by a slight decrease in identifications, which is proportional to the number of estimated targets per minute in the hybrid-DIA method (Supplementary Fig. 2C). This data could be used to predict the maximum limit of peptides that are realistic to target in a given gradient length by extrapolation (Supplementary Fig. 2D).

Finally, we evaluated the reproducibility between the quantification obtained from standard DIA and hybrid-DIA (Fig. 2E) and found positive correlation, comparable to replica runs in DIA (Supplementary Fig. 2E). Even so, the correlation decreased with lower input amounts, but this is most likely due to the higher variability introduced when doing phospho-enrichment with very low phosphopeptide enrichment inputs (>10 μg) (Fig. 2E).

### Benchmark of hybrid-DIA against SureQuant for targeted analysis of EGF stimulation

Having demonstrated the advantages of using hybrid-DIA compared to standard DIA runs, we next benchmarked its quantitative performance against the state-of-the-art spike-in triggered PRM acquisition method named SureQuant^10^ (Supplementary Fig. 1A). Phospho-enriched A549 digest dilution experiment (Fig. 2A) was acquired using both SureQuant and hybrid-DIA methods such that we could directly compare the sensitivity of the targeted scans in both approaches. We observed that both methodologies provide equivalent quantification performance through the dilution series range in terms of precision and accuracy with SureQuant equally affected by the relative abundance of the phosphopeptides (Fig. 3A).

**Figure 3.**
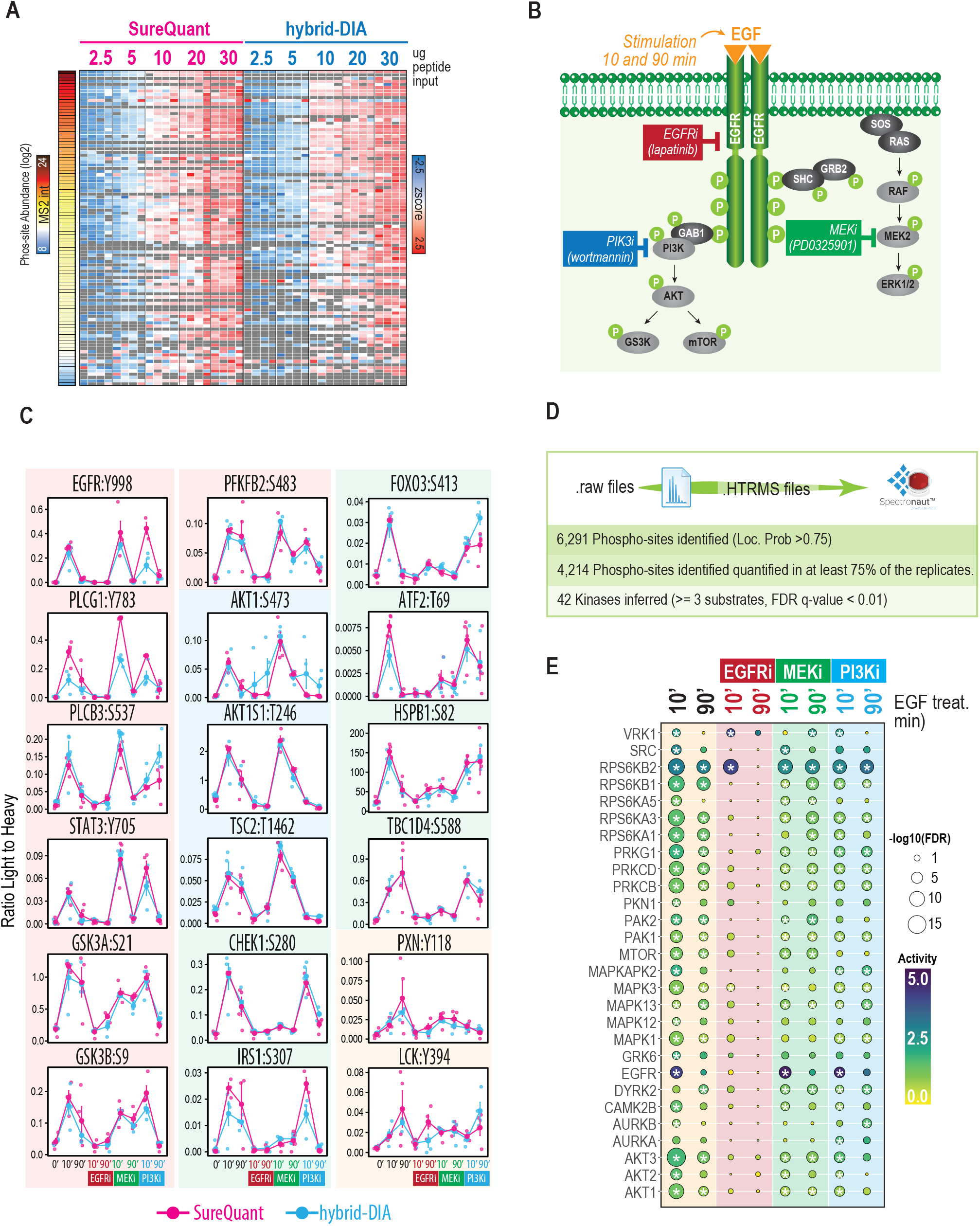
(A) Heatmap showing relative quantification of targeted peptides in a dilution series experiment (Fig 2A) in SureQuant (pink) and hybrid-DIA (blue). Sites are sorted by abundance reflecting that lowest abundant peptides show higher variability in both techniques. (B) Experimental design to study EGF time course stimulation in the presence of three inhibitors of downstream pathways: EGFRi, Pi3Ki and MEKi. Each condition was performed in quadruplicates. (C) Profile plot of absolute ratio Endogenous to Heavy standard of differentially regulated sites at 10 minutes of EGF stimulation in the hybrid-DIA dataset (two-sided t-test, BH-FDR). In blue, data from hybrid-DIA quantification (n=4); and in pink, data from SureQuant quantification (n=4). Background color is used to group the phospho-sites based on its response to each inhibitor: in red, sites that are targets of EGFR; in blue, sites that are downstream of PI3K and in green, sites that are downstream of MEK. (D) Summary of results obtained from the extracted DIA scans from the hybrid-DIA experiment. (E) Kinase activity inference analysis obtained from RoKAI using the discovery analysis data from the hybrid-DIA experiment. Asterisks indicate q-value < 0.01.

To extend this benchmark to a biological meaningful scenario, we performed EGF stimulation and chemical inhibition of downstream kinases in HeLa cells as a model of dynamic cellular signaling pathway rewiring (Fig. 3B). We selected this model system because the SureQuant Multipathway Phosphorylation Mix panel covers the EGFR signaling pathway, and downstream MEK and PIK3 kinase pathways. Moreover, we also scaled down the system to test both methods in most challenging conditions with limited input material growing cells in P6 plates to obtain approximately 50 μg of peptide per condition prior to phospho-enrichment. Our goal was to use the targeted data to reconstruct the phosphorylation pathway and infer the inhibited kinases. When we compared the quantitative profiles of the targeted peptides in SureQuant and hybrid-DIA of the sites differentially regulated by 10 minutes EGF stimulation, we observed that both methodologies provide equivalent results (Fig. 3C). Interestingly, with both SureQuant and hybrid-DIA and using the panel of synthetic heavy phosphopeptides, we can clearly identify kinase-specific responses with EGFR and AKT sites dynamically regulated by both EGF and kinase inhibitors, reflecting the potential of both methodologies to recapitulate kinase activity using this panel of peptides (Fig. 3C). Furthermore, the hybrid-DIA data also provided data related to the background phosphoproteome covering 6,163 sites (Fig. 3D), on top of the quantification of the sites from the panel. Using this data in a discovery phosphoproteomics pipeline, we could further reinforce the information on kinase inference (Fig. 3E). Using RoKAI^18^, we identified a rapid activation of multiple kinases upon EGF stimulation and how this activity was abrogated when inhibiting EGFR with Lapatinib (Fig. 3E). Furthermore, we observed that MTOR activity, which is a downstream target of PI3K, is specifically inhibited with Lapatinib (EGFRi) and wortmannin (PI3Ki), or that MAPK1 signaling, target of MEK, is significantly reduced after PD0325901 (MEKi) but not affected by wortmannin (PI3Ki) (Fig. 3E). Collectively, these results show the advantages of performing hybrid-DIA rather than only targeted acquisition methods as it maximizes the information retrieved from single-shot phosphoproteomics samples.

### Phosphoproteomics signature in 2D vs 3D model of colorectal cancer

Next, we decided to apply our intelligent data acquisition strategy to the most challenging biological in vitro models by studying dynamic phosphoproteome signaling in single multicellular cancer spheroids, and compare the signaling in these to conventional 2Dmonolayer culture of colorectal cancer cells. Three-dimensional tumor models, such as spheroids, offers an improved model to assess molecular and physiological aspects that are essential for drug development including drug penetration, hypoxic/necrotic environment, stemness and cell interaction, among many others^19,20^. Traditionally, spheroid models have been technically challenging, especially from a proteomics perspective, due to the low protein amount obtained from single spheroids, which typically requires pooling of several spheroids per condition to achieve a reasonable proteome coverage^21–23^. These limitations are even more evident when studying the phosphoproteome layer, due to the need for phospho-enrichment prior to LC-MS/MS measurements. Consequently, we reasoned that drug screening in single spheroids by phosphoproteomics was an ideal example of an experimental set-up requiring high sensitivity and benefitting from using our hybrid-DIA pipeline.

To achieve as good phosphopeptide coverage in single spheroids as possible, we have improved the sensitivity of our phosphoproteomics pipeline with the introduction of a modified phospho-enrichment protocol in combination with higher-resolution online chromatography to enhance MS sensitivity. For the latter, we took advantage of the higher sensitivity and chromatographic performance achieved with “whisper” nanoflow gradients on the Evosep One LC platform when using the Aurora column from IonOpticks (Supplementary Fig. 3A). Next, we observed that the use of MagReSyn^®^ ZrIMAC-HP beads^24^ outperformed MagReSyn^®^ TiIMAC-HP beads for low peptide input amounts in the low microgram range (Supplementary Fig. 3B). Additionally, we previously described how a second phosphopeptide-enrichment step in the Kingfisher platform is easily implemented by looping through the protocol, without the need to change buffers, but also reusing the beads and the elution buffer^25^. The implementation of the improved experimental protocol in combination with hybrid-DIA MS analysis, maximized the phospho-signaling information retrievable from single spheroids. We analyzed the dynamic phosphoproteome of single colon cancer spheroids in the context of chemotherapeutic treatment with 5-fluorouracil (5-FU), a common drug use in clinics for colorectal cancer treatment, and compared the results to the differential phosphoproteomics of 5-FU treated cells in monolayer cultures (Fig. 4A). For both monolayer and 3D culture, we seeded 20,000 cells per condition and grew them for three days until the spheroids were fully formed. At that time, 5-FU was added and samples collected at 0, 1, 3, 6, 12 and 24 hours after treatment with each condition as five independent replicates. Overall, the experiment consisted of 30 single spheroids and 30 samples of monolayer counterparts (Fig. 4A). All samples were lysed in 5% SDS, proteins extracted and trypsin digested using the protein aggregation capture (PAC) protocol and phosphopeptides enriched using Zr-IMAC HP beads. To each of the resulting 60 phosphopeptide samples, we spiked-in 50 fmol of the SureQuant MultiPathway Phosphorylation kit, which contained several cellular phosphorylation site markers of DNA damage and apoptosis, such as HSPB1:S82^26^, HSPB1:S15, JUN:S63^27^ and TP53:S3 1 5^28,29^. From the resulting raw MS files, we extracted the data from all targeted IS/ENDO MSx scans, and found 62 phosphorylation sites that were differentially regulated in at least one time point in either of the two conditions (3D-spheroids or 2D-monolayer) (Fig. 4B). Although the data from the targeted analysis revealed a similar response in terms of 5-FU activated signaling pathways in 2D monolayer cells and the 3D spheroid model, there were some notable differences. Substrate sites of the stress-responsive kinase, MAPKAPK-2 (MK2), HSPB1 Ser15 and Ser82 were phosphorylated at 6 to12 hours in monolayer culture, whilst their upregulation required up to 24 hours in spheroids (Fig. 4B-C). In contrast, apoptosis-activating phosphorylation sites on JUN Ser63 and TP53 Ser315 showed more synchronous temporal profiles in both systems, with significant activation as early as 3 hours for the Jun phosphorylation site (Fig. 4B-C). Interestingly, we also found phospho-sites related to MTOR and GSK3 signaling, which were specifically upregulated in spheroids only peaking at the latest 24 hours time point (Fig. 4B). The targeted analysis of this panel of phosphopeptides therefore serves as a highly sensitive and multiplexed assay to accurately probe the activity state and signaling dynamics of the major cellular kinase pathways directly informing about the signaling state of the cells analyzed. Furthermore, in addition to the phospho-signature extracted by the targeted quantification of the phosphopeptide panel, the hybrid-DIA MS data also contained comprehensive phosphoproteome profiling from the DIA scans. After conversion of the files to HTRMS format, we analyzed them with directDIA in Spectronaut (v17) obtaining quantification for 18,946 localized phospho-sites. To perform quantitative comparisons, we filtered the global phosphoproteomics dataset and retained 8,783 phospho-sites that were quantified in at least three out of five spheroids analyzed per one treatment time point, whilst 12,084 phospho-sites were quantified in the same proportion of samples in the monolayer culture condition (Fig. 5A). Such a phosphoproteome coverage is on par with data obtained in other large-scale phospho-proteomics screenings that use significantly higher peptide input material^5,25,30^. This reflects that the improvements in our sample preparation processing and MS analysis pipeline makes it realistic to scale down input amounts for highly sensitive phosphoproteomics experiments, such as analyzing single spheroid, while preserving a considerable coverage of the quantifiable phosphoproteome.

**Figure 4.**
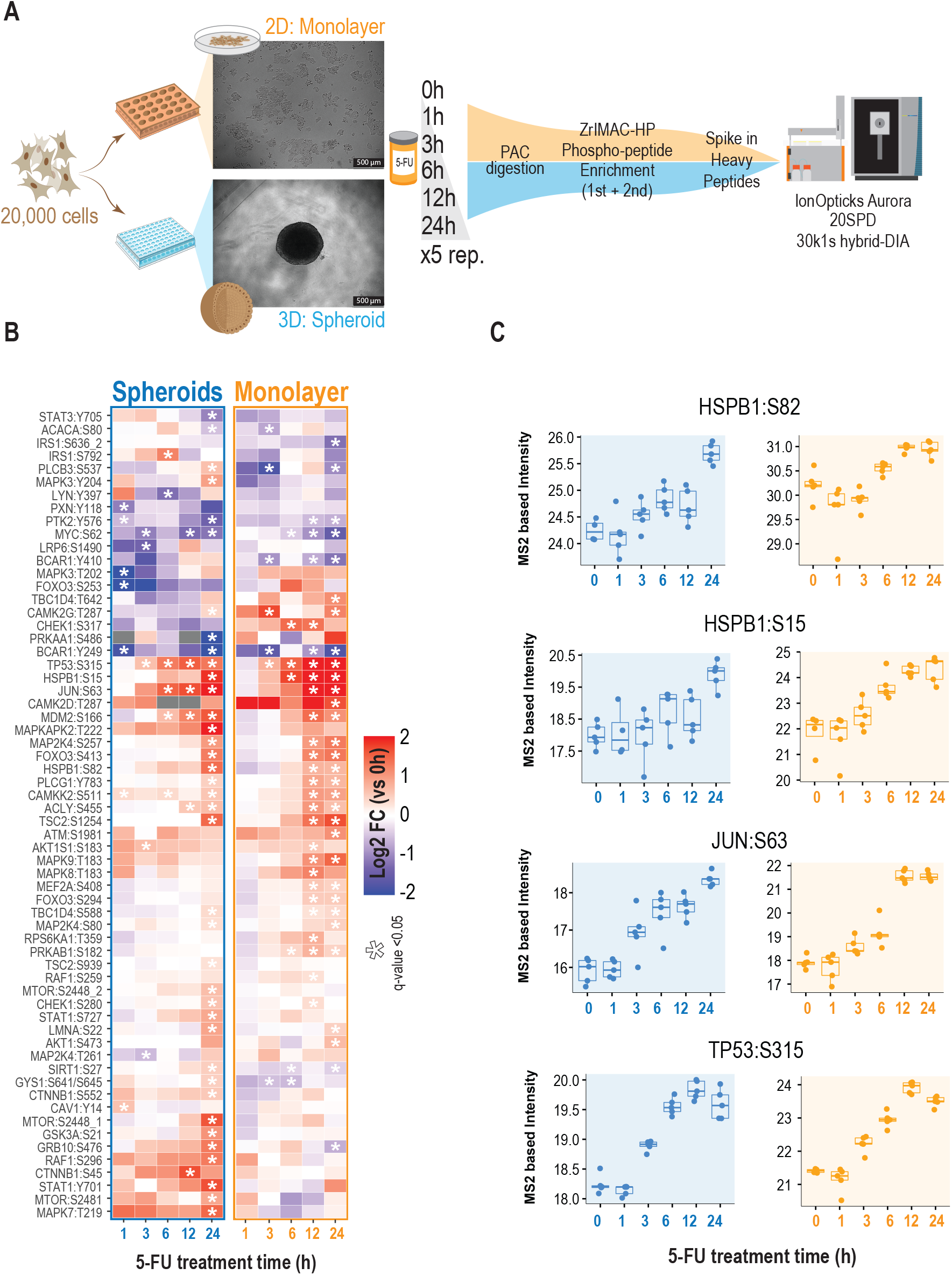
(A) Experimental design for the comparison of spheroids against monolayer culture of HCT116 cancer cells treated with 5-Fluorouracil. (B) Heatmap showing the phosphosites from the SureQuant™ Multipathway Phosphopeptide Standard panel that are differentially regulated (two-sided t-test, BH-FDR) in at least one point. Color indicates the average log2 fold change of each time point against time 0 (n=5). Asterisk indicates q-value <0.05. (C) Boxplot of MS2 intensities obtained from hybrid-DIA scans of relevant phosphorylation markers of DNA damage (n=5).

**Figure 5.**
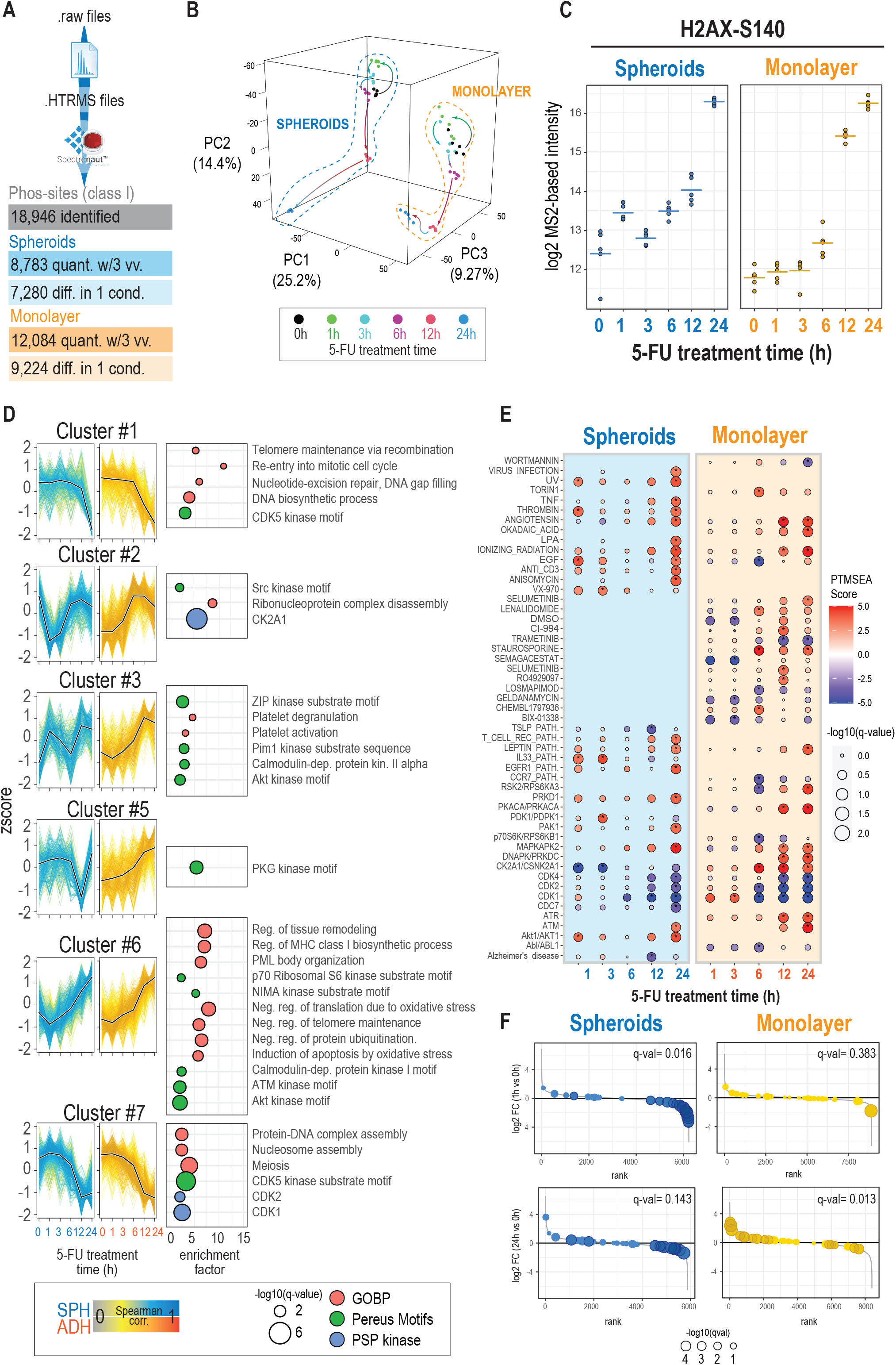
(A) Overview of the results obtained from the analysis of the DIA data with Spectronaut, after conversion to HTRMS format. (B) Principal Component analysis of spheroids and monolayer-grown cells treated with 5-FU at 0, 1, 3, 6, 12 and 24 hours (n=5). (C) Log2 intensity at MS2 level of H2AX Serine 140 (n=5, horizontal lines indicate the average of all measures). (D) Temporal profiles for relevant clusters (see Supplementary Figure 4A). To the right of each graph, the results from a Fisher’s exact test to show overrepresentation of terms from GOBP, Phosphositeplus Kinases and Kinase motifs. Size of the dot indicates the significance (two-sided, Fisher’s exact test, BH-FDR corrected), position on the x-axis, the enrichment factor, and the color indicates the ontology to which each term belongs. (E) PTMSEA results. Size of the dot indicates the significance (BH-FDR corrected); color indicates whether the term (associated pathway or kinase) is upregulated (red) or downregulated (blue). (F) Rank plots showing phospho-sites ranked by their fold change (log2) at 1 or 24 hours of treatment versus non-treated samples. Dots indicate the position of phospho-sites from the CK2A1 term (from PTMSEA database). Size of the dot indicates the significance of the fold change (limma robust moderated t-test, two-sided, BH-FDR, n=5). Darker dots highlight the sites with FDR corrected p-value < 0.01.

Principal Component Analysis (PCA) showed a clear separation between monolayer cells and spheroids in the first principal component (PC1), whereas the temporal effects of 5-FU followed similar trends in both conditions in principal components two and three (PC2 and PC3) (Fig. 5B). The 5-FU mechanism of action impairs DNA replication by inducing double-strand breaks (DBSs) during S phase of the cell cycle activating the DNA damage response^31–33^. Accordingly, we observed that phosphorylation of serine 140 in histone H2AX, a biomarker of DSBs known as gamma-H2AX, increases significantly upon treatment of colorectal cells with 5-FU (Fig. 5C). Interestingly, baseline levels of H2AX Ser140 were slightly higher in spheroids compared to adherent 2D monolayer-cultured cells, but, in contrast, monolayer-grown cells showed significantly higher increase in the phosphorylation site change of this marker when compared to spheroids, especially evident at 12 hours of treatment (Fig. 5C). This could indicate that monolayer-grown cells are more sensitive to the effect of 5-FU than spheroids, and highlights the importance of using 3D models exhibiting a different sensitivity to chemotherapeutic agents than cells grown in plates. The observed discrepancy in drug response kinetics is probably due to inherent delay in the diffusion of the drug into the inner spheroid in order to exert its full effect^34^. Furthermore, to identify 5-FU activated kinase signaling pathways in an unbiased manner, we performed an exploratory bioinformatic analysis of the full DIA phosphoproteomics dataset. We clustered the regulated phosphopeptide sites after z-scoring across the different drug-treatment time points and extracted the different temporal profiles observed in both treatments. This analysis revealed that the main biological response triggered by 5-FU are equivalent in both cell models: clusters #1 and #7 shows the downregulation of cell-cycle control by CDKs, and, conversely, cluster #6 shows a parallel upregulation of signaling pathways related to stress mediated by ATM and AKT kinases (Fig. 5D). However, this analysis also showed significant differences between the two cell models, for instance, the distinct temporal trend in the upregulation of AKT, or the specific early downregulation of CK2A1 in spheroids. We complemented this bioinformatic analysis using PTMSEA^35^ to annotate different phospho-regulated pathways and kinases in either 3D or 2D models across 5-FU treatment time. As expected, we found that MAPKAPK2 kinase, a stress-responsive kinase that has been connected with resistance in 5-FU treated colorectal cancer cells^36^, was upregulated by treatment with 5-FU. In line with the targeted phosphopeptide data, the activation of this kinase was evident earlier (12h) in the monolayer cells than in the spheroids (Fig. 5E). On the other hand, we found that cyclin dependent kinases 1, 2 and 4 activities were strongly downregulated after 5-FU treatment in both conditions but more significantly in monolayer cells (Fig. 5E). Interestingly, up-regulation of CDK1 and CDK4 is correlated with poor prognosis in cancer patients, especially in those with resistance to 5-FU^37–39^. Although most of the phospho-site signatures follow the same trend in spheroids and monolayer culture, we found casein kinase 2 alpha (CK2A1) to be strikingly different with opposite regulation (Fig 5D-E). There is evidence that CK2A1 levels correlate with poor prognosis in CRC patients^40^. Here we found a strong and rapid downregulation of this kinase in the spheroids, whilst it shows a slight upregulation in monolayer-grown cells (Fig 5F).

## Discussion

In this work, we present a MS acquisition method, termed hybrid-DIA, which enables intelligent acquisition of a predefined set of target peptides while simultaneously acquiring shotgun proteomics data using DIA. Consequently, our methodology combines accurate and sensitive quantification of targeted proteomics with the depth and unbiased discovery analysis of traditional DIA methods. Importantly, to facilitate the usage and analysis of data derived from using this method, we have designed a freely available data analysis pipeline.

We assume that this acquisition method is most beneficial for biomedical applications where sample input is limited and/or high-throughput is requested for large sample size analysis, and therefore critical to maximize the information that can be retrieved from a single-shot MS run. To serve as examples of such applications, we have performed highly sensitive phosphoproteomics of model systems with limited material using hybrid-DIA to prove the benefits of this workflow. As demonstrated by our data, hybrid-DIA benefits are more significant when targets of interest are of low abundance. This is because the targeted part of the method improves the limit of detection and quantification of predefined targets, which in standard DIA analysis will not be confidently detected.

One important consideration for the implementation of the hybrid-DIA workflow is that the inclusion of targeted scans in the DIA acquisition scheme is accompanied by prolonged MS acquisition cycle time, which needs to be carefully assessed when defining the number of targets. Ideally, the peptide targets must be evenly distributed across the chromatographic elution time range such that the number of targets per minute is regular through the run. Moreover, we presented guidelines on how to design the experiment based on the number of targets and the expected reduction in the depth of the proteome. For instance, in our set-up we estimated that 8 targets per minute would potentially lead to a reduction of 25% in overall DIA-based identifications (Supplementary Fig. 2D). Therefore, to prevent such losses it would be important to elongate the chromatographic gradient when scaling up the target list.

Most interestingly, not only have we demonstrated that both the DIA and targeted parts of the hybrid-DIA workflow are on par with state-of-art in proteomics for each method, but also the usefulness of hybrid-DIA methodology for clinical research purposes. Clinically relevant *in vitro* models in early drug screening are essential for developing potent and effective chemotherapeutic agents. Three-dimensional tumor models such as cancer cell spheroids more closely mimic in vivo solid tumors than monolayer cultured cell lines, and these 3D cell models have due to their similarity to tumor tissue in vivo in metabolic and proliferation gradient distribution emerged as attractive models for the early stages in drug screening^41,42^. However, due to the nature of spheroids, they are limited in size, and therefore in the amount of protein available for subsequent MS analysis^43^. Previous proteomics investigations relied on pooling of multiple single-spheroids for each condition analyzed^44^, which significantly reduces the throughput of this model. This limitation is even more relevant when studying the phosphoproteome, and to our knowledge, there is no prior art that provides comprehensive phosphoproteome profiles of single spheroids. Without doubt, a methodology that allows assessing phosphoproteomic response in single spheroids will truly increase the throughput of drug screening, and aid the field in the direction of using 3D models instead of 2D-monolayer grown cellular models.

## Code availability

Custom Python and R code used in the manuscript, both for the hybrid-DIA analysis, as well as for downstream data analysis, is available in the GitHub repository https://github.com/anamdv/HybridDIA. Hybrid-DIA API can be downloaded from https://github.com/thermofisherlsms.

PTM collapse plugin requires Perseus and R (minimum version 3.6.0) to run and it is available at https://github.com/AlexHgO/Perseus_Plugin_Peptide_Collapse.

## Author contribution

A.M-V. designed the experiments, optimized the API usage, performed all proteomics experiments, analyzed the data and implemented the data analysis pipeline. C.K. and G.F helped in experimental design and optimization of phosphorylation enrichment protocol. L.V.H. and G.F. generated the spheroids model. L.V.H. prepared the samples in the 5-FU treatment experiment. K.L.F, Y.X., T.M. and A.A.M. contributed to the development of the API. Y.C and Y.I. provided input on experiments and evaluated results. J.V.O. designed the experiments and critically evaluated the results. A.M.V., Y.X. and J.V.O. wrote the manuscript. All authors read, edited, and approved the final version of the manuscript.

## Acknowledgements

This project was in part funded through a research collaboration ‘cSHARP’ between Thermo Fisher Scientific, University of Copenhagen, University of Kyoto and Academia Sinica. Work at The Novo Nordisk Foundation Center for Protein Research (CPR) is funded in part by a generous donation from the Novo Nordisk Foundation (NNF14CC0001). This work has also been funded as part of EPIC-XS project under the grant agreement no. 823839 funded by the Horizon 2020 programme of the European Union and supported by the European Research Council through ERC-Synergy grant 810057-HighResCells. C.K. is supported by the Marie Skłodowska Curie European Training Network “PUSHH” (grant number No. 861389). We thank Aaron Gajadhar and Bhavin Patel from Thermo Fisher Scientific for providing us early access to the SureQuant™ Multipathway Phosphopeptide Standard kit.

## Online Methods

### Sample Prep: A549 dilution series for phosphoproteomics

A549 (ATCC CCL-185) were cultured in DMEM (Gibco, Invitrogen), supplemented with 10% fetal bovine serum (FBS, Gibco), 100U/ml penicillin (Invitrogen), 100 μg/ml streptomycin (Invitrogen), at 37 °C, in a humidified incubator with 5% CO2. Cells were harvested at ~80% confluence by washing twice with PBS (Gibco, Life technologies) and subsequently adding boiling lysis buffer (5% sodium dodecyl sulfate (SDS), 5 mM tris(2-carboxyethyl)phosphine (TCEP), 10 mM chloroacetamide (CAA), 100 mM Tris, pH 8.5) directly to the plate. The cell lysate was collected by scraping the plate and boiled for an additional 10 min followed by micro tip probe sonication (Vibra-Cell VCX130, Sonics, Newtown, CT) for 2 min with pulses of 1 second on and 1 second off at 80% amplitude. Protein concentration was estimated by BCA. Protein was digested using the Protein Aggregation Capture protocol. Briefly 1 mg of protein was resuspended with acetonitrile to a final 70% concentration. MagReSyn^®^ Hydroxyl beads were added in a proportion 1:2 (protein:beads). Protein aggregation was performed in two steps of 1 minute mixing at 1000 rpm, followed by 10 minute pause each. Beads were subsequently washed three times with 1 ml 95% ACN and two times with 1ml 70% EtOH. 300 μl of digestion buffer (50mM Ammonium Bicarbonate) and proteases were added in the following proportions: trypsin 1:250 (enzyme:protein) and lysC 1:500 (enzyme:protein). Digestion was carried out overnight at 37 °C with looping mixing. Protease activity was quenched by acidification with trifluoroacetic acid (TFA) to a final concentration of 1%, and the resulting peptide mixture was concentrated on Sep-Pak (C18 Classic Cartridge, Waters, Milford, MA). Peptides were eluted with 150 μl 40% ACN, followed by 150 μl 60% ACN. The combined eluate was reduced by SpeedVac (Eppendorf, Germany) and the final peptide concentration was estimated by measuring absorbance at 280 nm on a NanoDrop 2000C spectrophotometer (Thermo Fisher Scientific). For phosphoproteomic enrichment, each peptide amount (30, 20, 10, 5 and 2.5 μg) were resuspended with 200 μl of Loading Buffer (80% ACN; 5%TFA, 1M Glycolic Acid). Subsequent phospho-enrichment was performed in the King-fisher robot using 5 μl of MagReSyn^®^ Ti-IMAC HP beads (20mg/ml) ^13^. Enriched phosphopeptides were acidified with 10% TFA until pH<3 and filtered to remove in-suspension particles (1 min, 500 g, MultiScreenHTS HV Filter Plate, 0.45 μm, clear, non-sterile). 0.5 μl of the SureQuant™ Multipathway Phosphopeptide Standard (100 fmol/μL) was added to each sample prior loading into Evotips for subsequent MS analysis.

### Sample Prep: HeLa stimulation with EGF and kinase inhibitors

HeLa (ATCC CCL-2) cells were grown in a P6 dish until 70% confluence. Cells were serum-starved for 6 hours. Control HeLa cells were stimulated with 100 ng/mL of EGF for 10 and 90 minutes. For the drug inhibitor treatment, cells were initially incubated in each inhibitor (Lapatinib 14μM, PD0325901 3μM and Wormannin 25μM) for 15 minutes. Then cell were stimulated with 100 ng/mL of EGF for 10 and 90 minutes, in the presence of the inhibitors. Cells were lysed with 200 μl of boiling lysis buffer (5% SDS; 100mM Tris pH 8.5, 5mM TCEP and 10mM CAA) and incubated at 95 °C, for 10 minutes with mixing (1000 rpm). Lysates were sonicated with a 8-tip probe (1 minute, 1 second ON, 1 second OFF, 50% amplitude, 8-channel Fisherbrand™ Tip Horn for Model 120 Sonic Dismembrator). Protein concentration was calculated by BCA. 150 μg of protein was digested using the Protein Aggregation Capture protocol in the Kingfisher Robot as detailed above. Digested peptides were acidified after digestion with TFA to final 1% concentration and loaded into a Sep-Pak tC18 96-well Plate, (40 mg Sorbent per Well, Waters) for desalting. Peptides were eluted in 75ul of 80% ACN and resuspended with 150 μl of Concentrated Loading Buffer (80% ACN; 8%TFA, 1.6M Glycolic Acid). 0.5 μl of the SureQuant™ Multipathway Phosphopeptide Standard (100 fmol/μL) was added to each sample and continued for subsequent phospho-enrichment in the Kingfisher robot using 5ul of MagReSyn^®^ Ti-IMAC HP beads (20mg/ml). Enriched phosphopeptides were acidified with 10% TFA until pH<3 and filtered to remove in-suspension particles (1 min, 500 g, MultiScreenHTS HV Filter Plate, 0.45 μm, clear, non-sterile). Finally, samples were loaded into Evotips for subsequent MS analysis.

### Sample Prep: Sensitive phosphoproteomics on Single Spheroids and Monolayer culture of HCT116 cancer cells treated with 5-Fluorouracil

Multicellular spheroids and monolayer culture cells were grown from HCT116 cells (ATCC CCL-247). Prior to seeding, cells were harvested from normal cell plates and counted. For spheroids generation, 20,000 cells were seeded on ultra-low attachment 96-well plates (Corning CoStar, Merck) and cultured in 90% DMEM (Gibco, Invitrogen), supplemented with 10% heat-inactivated fetal bovine serum (FBS, Gibco) and 10,000 U/mL penicillin and streptomycin (Invitrogen). Subsequently, spheroids were cultured for 96 hours hours at 37 °C, in a humidified incubator with 5% CO2. Cell medium was refreshed after 48 hours, by aspirating half the old medium (making sure not to alter the spheroid) and adding the same amount of fresh medium. For monolayer culture, 20,000 cells were seeded on 24-well plates. After 96 hours the spheroids and monolayer cultured cells were treated with 1.8 μM 5-fluorouracil (Sigma-Aldrich) for 1, 3, 6, 12 and 24 hours. Subsequently the spheroids were harvested by resuspension in 200 μl boiling lysis buffer (5% SDS, 5mM TCEP, 10mM CAA, 100mM T ris pH 8.5) and mixed in a thermo-shaker (1000 rpm) at 95 °C until the entire spheroid disaggregates (approximately 10 minutes). Monolayer cultured cells were lysed with 200 μl of boiling lysis buffer (5% SDS; 100mM Tris pH 8.5, 5mM TCEP and 10mM CAA) and incubated at 95 °C, for 10 minutes with mixing (1000 rpm). Afterwards, lysates were sonicated with a probe (1 minute, 1 second ON, 1 second OFF, 50% amplitude, 2mm Fisherbrand™ Probe for Model 120 Sonic Dismembrator).

Lysates were digested using the Protein Aggregation Capture protocol in the Kingfisher Robot modified for low input amounts. The ratio of MagReSyn^®^ Hydroxyl beads to protein used was 16:1, and the ratio of enzymes used was 1:100 for lysC and 1:50 for trypsin. Samples were digested for 6 hours in 200 μl of 50 mM triethylammonium bicarbonate. Digested peptides were acidified after digestion with 50 μl of 10% formic acid. Peptides were concentrated in a SpeedVac at 45 °C until volume was 20μl. Peptides were resuspended in Loading Buffer (80% ACN; 5%TFA, 0.1 M Glycolic Acid) and subjected to phospho-enrichment in the Kingfisher Robot. 5 ul of MagReSyn^®^ ZrIMAC-HP beads (20mg/ml) were used per sample, and two sequential enrichment were performed, without changing buffers in between. Samples were eluted in 200 μl of 1 % NH3OH and subsequently acidified with 40 μl of 10 %TFA. Prior to evotipping, samples were filtered (1 min, 500 g, MultiScreenHTS HV Filter Plate, 0.45 μm, clear, non-sterile). 0.5 μl of the SureQuant™ Multipathway Phosphopeptide Standard (100 fmol/μL) was added to each sample. Finally, samples were loaded into Evotips for subsequent MS analysis.

### Implementation of hybrid-DIA scans on a quadrupole Orbitrap mass spectrometer

The Orbitrap Exploris 480 mass spectrometer was operated with the instrument control software Tune v3.0 or higher (Thermo Fisher Scientific). A standard DIA MS method was built within Xcalibur (v4.3, Thermo Fisher Scientific). Hybrid-DIA scans were customized and programmed via an application interfacing program (API) tool (v1.3 or higher) provided by Thermo Fisher Scientific. A full guide on how to operate the API is provided as Supplementary Data 1. Briefly, the hybrid-DIA API works as follows:

1. The theoretical mass-to-charge value, charge state, and retention time window of internal standard (IS) peptides and corresponding endogenous (ENDO) peptides, as well as the theoretical mass-to-charge values of the fragments of IS peptides are predefined as an input .txt file for hybrid-DIA API program. Moreover, the following parameters for the MS2 acquisition are indicated in the API graphic interface: acquisition time, mass tolerance, defined first mass, NCE, isolation width, AGC target, maximum injection time in milliseconds, MS Trigger Intensity Threshold and dynamic exclusion (in seconds).
2. The precursors of IS peptides are analyzed in MS scans. When IS peptides are detected within the given retention time range, predefined mass tolerance, and above the intensity threshold in MS scan, a fast multiplexed (MSx) PRM MS/MS scan of all detected IS peptides is inserted and performed.
3. When a threshold of predefined fragments for any IS peptide are detected in the PRM MS2 scans within the defined mass tolerance, a multiplexed PRM MS2 scan of the IS peptide and its corresponding endogenous peptide (ENDO) is performed, where the maximal ion injection times for IS and endogenous peptide are set individually to maximize the detection sensitivity for the low abundant endogenous peptide while maintaining a fast DIA cycle time. These co-isolation scans occur for entire list of successfully analyzed IS peptides.
4. If step #2 fails to match the predefined conditions, the mass spectrometry continuously acquires the standard DIA data.
5. Following the completion of all MSx PRM scans of IS peptides and their corresponding ENDO peptide pairs (step #3), the mass spectrometry continuously acquires the standard DIA data. Steps #2 and #3 will repeat whenever the predefined precursors and fragments are identified, respectively.

### LC-MS/MS Analysis

Samples were analyzed on the Evosep One system using EV-1112 column (PepSep, 15 cm x 75um, beads 1.9 um) and EV-1087 emitter (Fused silica, 20μm). The column temperature was maintained at 40 °C using a butterfly heater (PST-ES-BPH-20, Phoenix S&T) and interfaced online using an EASY-Spray™ source with the Orbitrap Exploris 480 MS (Thermo Fisher Scientific, Bremen, Germany) using Xcalibur (tune version 3.0 or higher). Alternatively, the 5-fluorouracil treated samples were analyzed using an IonOpticks Aurora™ column (15cm-75um-C18 1.6um) interfaced with the Orbitrap Exploris 480 MS using a Nanospray Flex™ Ion Source with an integrated column oven (PRSO-V1, Sonation, Biberach, Germany) to maintain the temperature to 50 °C. In all samples, spray voltage was set to 1.8 kV, funnel RF level at 40, and heated capillary temperature at 275 °C. All experiments were acquired using 20 samples per day (SPD) gradient, except for the target dilution and phospho-optimization experiments, which were acquired using 40 SPD.

For full phospho-proteome hybrid-DIA analysis, full MS resolution were set to 120,000 at m/z 200 and full MS AGC target was 300% with an IT of 45 ms. Mass range was set to 350 - 1400. AGC target value for DIA scans was set at 1000%. Resolution was set to 30,000 and IT to 54 ms and normalized collision energy was 27%. DIA windows scanning from 472 to 1143 m/z with 1 m/z overlap were used (i.e. 11 windows of 61.1 Da for 1 second cycle time at 30K resolution). To enable non-isochronous injection times for MSx scans, the options must be enabled in Tune (available in Diagnostics > Method Setup).

For hybrid-DIA inclusion lists, the retention time schedule was calculated from Survey Scans runs, where an inclusion list containing the m/z and charge of the spiked-in IS peptides was used to specifically trigger their acquisition. In particular, for the A549 dilution series experiment, as well as for the EGF+Inhibitors experiment, the retention time schedule was obtained from the SureQuant runs used in those experiments. In both cases, data was imported to Skyline, where the peptide peak integration was manually validated, and the retention times were exported for hybrid-DIA analysis.

For SureQuant acquisition, we used the template available in Thermo Orbitrap Exploris Series Method Editor. Full-scan mass spectra were collected with a scan range: 300–1,500 m/z, AGC target value: 300%, maximum IT of 50 ms and 120,000 resolution. Several branches were used, each one for a unique isotopically labeled amino acid and charge state, which will determine the m/z offset. In particular, the method contained 8 branches for +2, +3 and +4 charge states of IS lysine (K8+) and arginine (R10+), as well as +3 charge state of IS alanine (A4+) and +2 charge state of IS valine (V6+) peptides. In each branch, the peptide m/z, charge and intensity thresholds are defined in the “Targeted Mass” filter node. For all peptides, intensity threshold was fixed to 1e5. Next, parameters for the “fast/survey” ddMS2 scans are defined. Resolution was set to 7,500 and IT to 10 ms and normalized collision energy was 27%. This is followed by the “Targeted Mass Trigger” filter node, which defines up to 6 product ions used for pseudo-spectral matching, allowing 10 ppm mass tolerance and minimum of at least 3 detected fragments for each precursor. This step is followed by a “sensitive/triggered” ddMS2 scan. For the sensitive scan, we used a specific isolation offset for each branch. Resolution was set to 60,000 and IT to 116 ms and normalized collision energy was 27%.

In both acquisition methods, inclusion list for peptides contained in the SureQuant™ Multipathway Phosphorylation Mix was reduced from 131 to 129, due to lack of detection of the precursors of two peptides from that mix (i.e. GSK3:S9 and TSC2:S1387).

### Data analysis: DIA-based Discovery Pipeline

For hybrid-DIA analysis, DIA scans were extracted using the HTRMS convertor tool from Spectronaut (v15.4 or higher) indicating hybrid-DIA conversion in “Conversion type”. HTRMS resulting files were further used for directDIA search in Spectronaut (v17).

MS files, both from standard DIA (raw) and hybrid-DIA (HTRMS) were searched using Spectronaut with a library-free approach (directDIA) using a human database (Uniprot reference proteome 2022 release, 20,598 entries). Carbamidomethylation of cysteines was set as a fixed modification, whereas oxidation of methionine, acetylation of protein N-termini and phosphorylation of serine, threonine and tyrosine were set as possible variable modifications. In the HeLa+EGF and Spheroids experiment, we filtered out ‘b-ions’ to prevent quantitative interference from Heavy peptides. The maximum number of variable modifications per peptide was limited to 5. PTM localization Filter was checked and PTM localization cutoff was set to 0.75. Cross-run normalization was turned off.

Phosphopeptide quantification data was exported and collapsed to site information using the plugin described in Bekker-Jensen et al^5^ (see Code Availability) in Perseus (v1.6.5.0). Phospho-sites intensities were log2 transformed and values were filtered to keep only phospho-sites quantified in at least 3 replicates in one experimental condition. Data was exported and further processed in R (v4.1.1). Normalization was performed using loess function from limma package (v3.50.3)^45^. Imputation of missing values was performed in two steps using the DAPAR package (v1.26.1)^46^ taking into account the nature of the missing values, as described by Lazar et al^47^ First, we considered partially observed values as those values missing within a condition in which there are valid quantitative values in other replicates. These partially observed values were imputed using the “slsa” function. Secondly, values missing in an entire condition were imputed using the detQuant function from imp4p package (v1.2). Finally, differential expressed phosphor-sites were calculated using limma (two-sided, BH FDR < 5%, robust), requiring at least three valid values in one of the two experimental conditions compared.

### Data analysis: Targeted Pipeline

Raw files acquired in hybrid-DIA mode were processed to extract separately the DIA scans for full phosphoproteome analysis and the IS/ENDO multiplexed scans for targeted analysis. Multiplexed scans containing the internal standard and the endogenous peptide were extracted in an mzML file using an in-house designed python GUI that combines the python library pymsfilereader (https://github.com/frallain/pymsfilereader) and MSConvert^48,49^. Resulting mzML files were loaded into a Skyline-daily (21.1.9.353) to extract the intensity information of IS/ENDO scans. Resulting files were used for injection time correction and peak area (AUC) calculation using the R-based shiny-app developed for this purpose. A complete guide to further process the hybrid-DIA scans and perform the IT normalization is available as Supplementary Data 2. The python GUI as well as the shiny-app for IT normalization and visualization are available as the github page for this project: https://github.com/anamdv/HybridDIA.

Skyline template containing the phosphopeptide library of the SureQuant Multipathway Phosphorylation Kit was initially provided by Thermo Fisher Scientific, but then manually curated per experiment to remove shared fragments between isoforms and interfering transients. SureQuant raw files and mzML files from hybrid-DIA runs were imported into Skyline using the abovementioned template using specific transition settings for each acquisition method (Table 1).

**Table 1.**
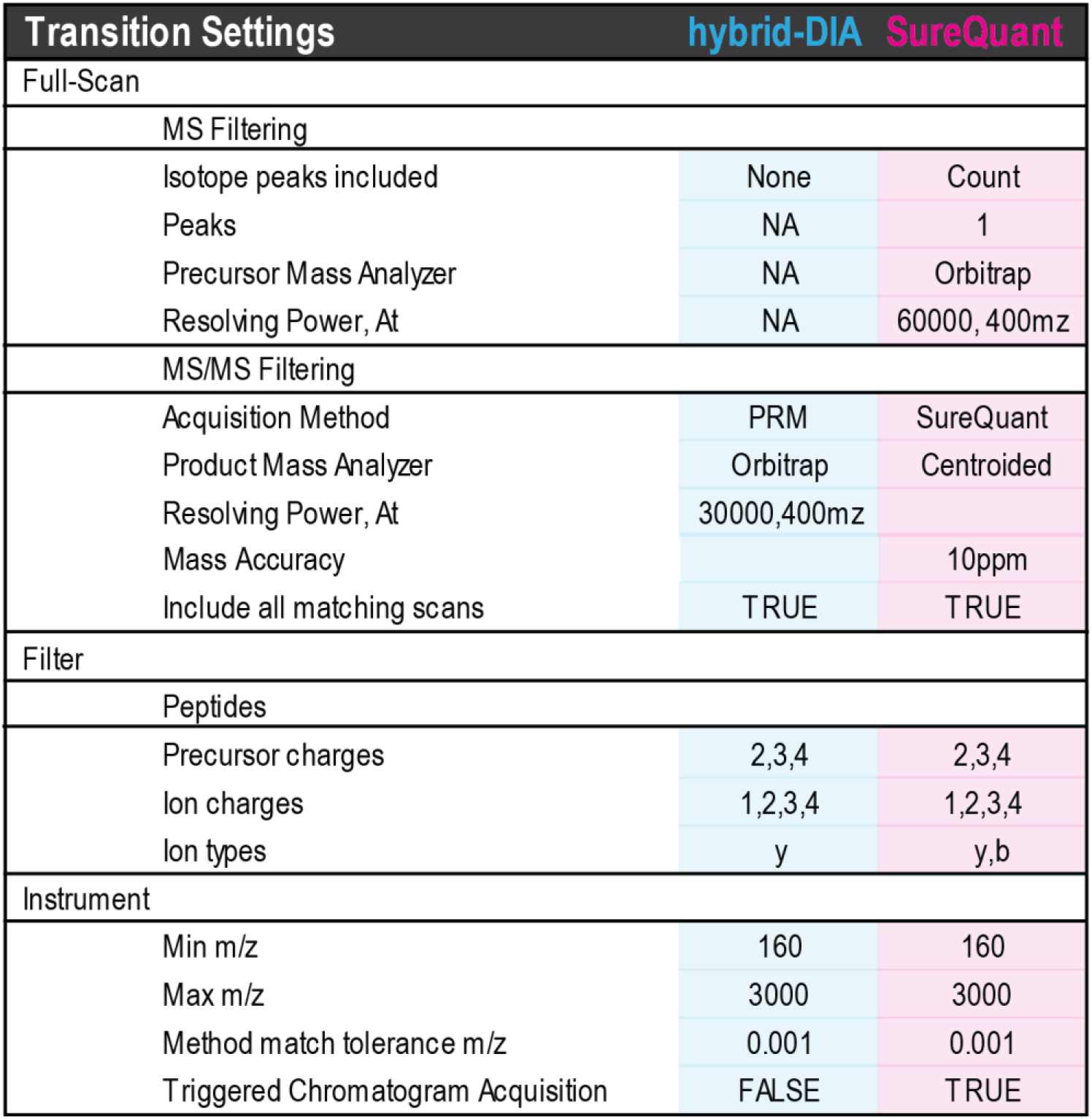
Transition Settings used in Skyline when importing hybrid-DIA or SureQuant data.

Hybrid-DIA quantification was performed using the AUC calculated as mentioned above. Intensities from ENDO peptides were normalized based on the IS peptide intensities. If a significant bias on peptide loading was observed in the DIA data (such in the spheroid dataset), a second normalization step was performed, using median intensity from DIA scans to correct ENDO peptide intensity.

SureQuant quantification was extracted directly from Skyline, using the data from the “Quantification_IS-ENDO” report, in particular from the column “Ratio To Standard”.

## References

1. Ludwig, C. et al. Data-independent acquisition-based SWATH-MS for quantitative proteomics: a tutorial. Mol. Syst. Biol. 14, e8126 (2018).

2. Ye, Z., Mao, Y., Clausen, H. & Vakhrushev, S. Y. Glyco-DIA: a method for quantitative O-glycoproteomics with in silico-boosted glycopeptide libraries. Nat. Methods 16, 902–910 (2019).

3. Robinson, A. E. et al. Lysine and Arginine Protein Post-translational Modifications by Enhanced DIA Libraries: Quantification in Murine Liver Disease. J. Proteome Res. 19, 4163–4178 (2020).

4. Tanzer, M. C., Bludau, I., Stafford, C. A., Hornung, V. & Mann, M. Phosphoproteome profiling uncovers a key role for CDKs in TNF signaling. Nat. Commun. 12, 6053 (2021).

5. Bekker-Jensen, D. B. et al. Rapid and site-specific deep phosphoproteome profiling by data-independent acquisition without the need for spectral libraries. Nat. Commun. 11, 1–12 (2020).

6. Lou, R. et al. DeepPhospho accelerates DIA phosphoproteome profiling through in silico library generation. Nat. Commun. 12, 6685 (2021).

7. Batth, T. S., Francavilla, C. & Olsen, J. V. Off-line high-pH reversed-phase fractionation for in-depth phosphoproteomics. J. Proteome Res. 13, 6176–6186 (2014).

8. Humphrey, S. J., Karayel, O., James, D. E. & Mann, M. High-throughput and high-sensitivity phosphoproteomics with the EasyPhos platform. Nat. Protoc. 13, 1897–1916 (2018).

9. Post, H. et al. Robust, Sensitive, and Automated Phosphopeptide Enrichment Optimized for Low Sample Amounts Applied to Primary Hippocampal Neurons. J. Proteome Res. 16, 728–737 (2017).

10. Stopfer, L. E. et al. High-Density, Targeted Monitoring of Tyrosine Phosphorylation Reveals Activated Signaling Networks in Human Tumors. Cancer Res. 81, 2495–2509 (2021).

11. Erickson, B. K. et al. A Strategy to Combine Sample Multiplexing with Targeted Proteomics Assays for High-Throughput Protein Signature Characterization. Mol. Cell 65, 361–370 (2017).

12. Grossegesse, M., Hartkopf, F., Nitsche, A. & Doellinger, J. Stable Isotope-Triggered Offset Fragmentation Allows Massively Multiplexed Target Profiling on Quadrupole-Orbitrap Mass Spectrometers. J. Proteome Res. 19, 2854–2862 (2020).

13. Bekker-Jensen, D. B. et al. A compact quadrupole-orbitrap mass spectrometer with FAIMS interface improves proteome coverage in short LC gradients. Mol. Cell. Proteomics 19, 716–729 (2020).

14. Deutsch, E. W. Mass spectrometer output file format mzML. Methods Mol. Biol. 604, 319–31 (2010).

15. Pino, L. K. et al. The Skyline ecosystem: Informatics for quantitative mass spectrometry proteomics. Mass Spectrom. Rev. 39, 229–244 (2020).

16. Chiva, C. et al. Isotopologue Multipoint Calibration for Proteomics Biomarker Quantification in Clinical Practice. Anal. Chem. 91, 4934–4938 (2019).

17. Ochoa, D. et al. The functional landscape of the human phosphoproteome. Nat. Biotechnol. 38, 365–373 (2020).

18. Yilmaz, S. et al. Robust inference of kinase activity using functional networks. Nat. Commun. 12, 1177 (2021).

19. Karlsson, H. et al. A novel tumor spheroid model identifies selective enhancement of radiation by an inhibitor of oxidative phosphorylation. Oncotarget 10, 5372–5382 (2019).

20. Karlsson, H., Fryknäs, M., Larsson, R. & Nygren, P. Loss of cancer drug activity in colon cancer HCT-116 cells during spheroid formation in a new 3-D spheroid cell culture system. Exp. Cell Res. 318, 1577–85 (2012).

21. Wang, L.-T., Proulx, M.-È., Kim, A. D., Lelarge, V. & McCaffrey, L. A proximity proteomics screen in three-dimensional spheroid cultures identifies novel regulators of lumen formation. Sci. Rep. 11, 22807 (2021).

22. Abe, Y. et al. Improved phosphoproteomic analysis for phosphosignaling and active-kinome profiling in Matrigel-embedded spheroids and patient-derived organoids. Sci. Rep. 8, 11401 (2018).

23. Steinmetz, J. et al. Descriptive Proteome Analysis to Investigate Context-Dependent Treatment Responses to OXPHOS Inhibition in Colon Carcinoma Cells Grown as Monolayer and Multicellular Tumor Spheroids. ACS omega 5, 17242–17254 (2020).

24. Arribas Diez, I. et al. Zirconium(IV)-IMAC Revisited: Improved Performance and Phosphoproteome Coverage by Magnetic Microparticles for Phosphopeptide Affinity Enrichment. J. Proteome Res. 20, 453–462 (2021).

25. Koenig, C., Martinez-Val, A., Franciosa, G. & Olsen, J. V. Optimal analytical strategies for sensitive and quantitative phosphoproteomics using TMT-based multiplexing. Proteomics 22, e2100245 (2022).

26. Rane, M. J. et al. Heat shock protein 27 controls apoptosis by regulating Akt activation. J. Biol. Chem. 278, 27828–35 (2003).

27. Li, L., Feng, Z. & Porter, A. G. JNK-dependent phosphorylation of c-Jun on serine 63 mediates nitric oxide-induced apoptosis of neuroblastoma cells. J. Biol. Chem. 279, 4058–65 (2004).

28. Wang, Y. & Prives, C. Increased and altered DNA binding of human p53 by S and G2/M but not G1 cyclin-dependent kinases. Nature 376, 88–91 (1995).

29. Abraham, J., Kelly, J., Thibault, P. & Benchimol, S. Post-translational modification of p53 protein in response to ionizing radiation analyzed by mass spectrometry. J. Mol. Biol. 295, 853–64 (2000).

30. Kitata, R. B. et al. A data-independent acquisition-based global phosphoproteomics system enables deep profiling. Nat. Commun. 12, 2539 (2021).

31. Wyatt, M. D. & Wilson, D. M. Participation of DNA repair in the response to 5-fluorouracil. Cell. Mol. Life Sci. 66, 788–99 (2009).

32. Srinivas, U. S. et al. 5-Fluorouracil sensitizes colorectal tumor cells towards double stranded DNA breaks by interfering with homologous recombination repair. Oncotarget 6, 12574–86 (2015).

33. Li, L. S. et al. DNA mismatch repair (MMR)-dependent 5-fluorouracil cytotoxicity and the potential for new therapeutic targets. Br. J. Pharmacol. 158, 679–92 (2009).

34. LaBonia, G. J., Lockwood, S. Y., Heller, A. A., Spence, D. M. & Hummon, A. B. Drug penetration and metabolism in 3D cell cultures treated in a 3D printed fluidic device: assessment of irinotecan via MALDI imaging mass spectrometry. Proteomics 16, 1814–21 (2016).

35. Krug, K. et al. A Curated Resource for Phosphosite-specific Signature Analysis. Mol. Cell. Proteomics 18, 576–593 (2019).

36. Yang, S. Y. et al. Inhibition of the p38 MAPK pathway sensitises human colon cancer cells to 5-fluorouracil treatment. Int. J. Oncol. 38, 1695–702 (2011).

37. Ding, X., Duan, H. & Luo, H. Identification of Core Gene Expression Signature and Key Pathways in Colorectal Cancer. Front. Genet. 11, 45 (2020).

38. Zhao, P., Hu, Y.-C. & Talbot, I. C. Expressing patterns of p16 and CDK4 correlated to prognosis in colorectal carcinoma. World J. Gastroenterol. 9, 2202–6 (2003).

39. Sung, W.-W. et al. High nuclear/cytoplasmic ratio of Cdk1 expression predicts poor prognosis in colorectal cancer patients. BMC Cancer 14, 951 (2014).

40. Lin, K.-Y. et al. Overexpression of nuclear protein kinase CK2 α catalytic subunit (CK2α) as a poor prognosticator in human colorectal cancer. PLoS One 6, e17193 (2011).

41. Breslin, S. & O’Driscoll, L. Three-dimensional cell culture: the missing link in drug discovery. Drug Discov. Today 18, 240–9 (2013).

42. Yue, X., Lukowski, J. K., Weaver, E. M., Skube, S. B. & Hummon, A. B. Quantitative Proteomic and Phosphoproteomic Comparison of 2D and 3D Colon Cancer Cell Culture Models. J. Proteome Res. 15, 4265–4276 (2016).

43. Feist, P. E., Sun, L., Liu, X., Dovichi, N. J. & Hummon, A. B. Bottom-up proteomic analysis of single HCT 116 colon carcinoma multicellular spheroids. Rapid Commun. Mass Spectrom. 29, 654–8 (2015).

44. Beller, N. C., Lukowski, J. K., Ludwig, K. R. & Hummon, A. B. Spatial Stable Isotopic Labeling by Amino Acids in Cell Culture: Pulse-Chase Labeling of Three-Dimensional Multicellular Spheroids for Global Proteome Analysis. Anal. Chem. 93, 15990–15999 (2021).

45. Ritchie, M. E. et al. limma powers differential expression analyses for RNA-sequencing and microarray studies. Nucleic Acids Res. 43, e47–e47 (2015).

46. Wieczorek, S. et al. DAPAR & ProStaR: software to perform statistical analyses in quantitative discovery proteomics. Bioinformatics 33, 135–136 (2017).

47. Lazar, C., Gatto, L., Ferro, M., Bruley, C. & Burger, T. Accounting for the Multiple Natures of Missing Values in Label-Free Quantitative Proteomics Data Sets to Compare Imputation Strategies. J. Proteome Res. 15, 1116–1125 (2016).

48. Chambers, M. C. et al. A cross-platform toolkit for mass spectrometry and proteomics. Nat. Biotechnol. 30, 918–20 (2012).

49. Holman, J. D., Tabb, D. L. & Mallick, P. Employing ProteoWizard to Convert Raw Mass Spectrometry Data. Curr. Protoc. Bioinforma. 46, 13.24.1–9 (2014).

