## Supplementary Data for "Hybrid-DIA: Intelligent Data Acquisition for Simultaneous Targeted and Discovery Phosphoproteomics in Single Spheroids"

- Supplementary Data 1: instructions to use the hybrid-DIA API in and Exploris 480 MS.
- Supplementary Data 2: guide for post-acquisition processing analysis of targeted scans from hybrid-DIA runs.
- Supplementary Figures 1-4

#### **Supplementary Data 1.** Instructions to use the hybrid-DIA API in and Exploris 480 MS.

In order to use hybrid-DIA acquisition in Exploris 480, follow these steps.

1. Download the folder "Exploris" from https://github.com/thermofisherlsms/MoonshotApps onto your local computer. Make sure the following items are present: Moonshot_V1_4.exe Thermo.API.Exploris-1.0.dll, Thermo.API.Spectrum-1.0.dll and Thermo.API-2.0.dll.
2. Ensure that your Exploris Instrument has an API license.
3. Double click the application "Moonshot_V1.4.exe" within the "Exploris" folder. The GUI shown in Figure 1 should open and the console message box shown in Figure 2.

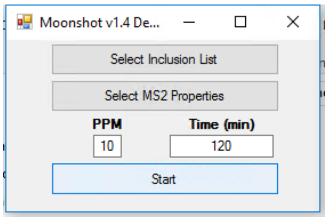


*Figure 1. Main screen of the Moonshot API Interface*

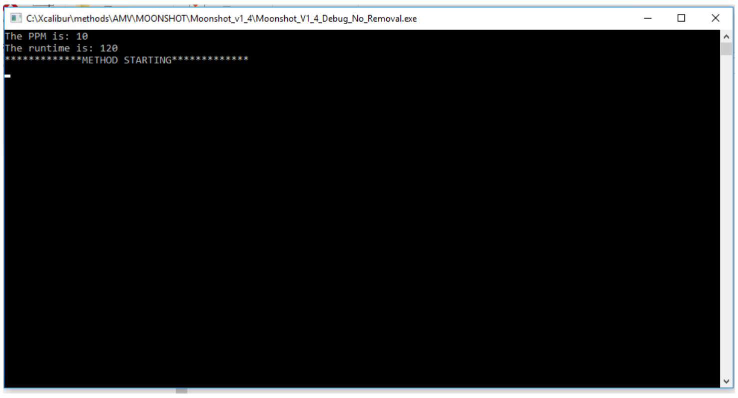


*Figure 2. Secondary console window. Messages from the program will be displayed here.*

4. Press "Select Inclusion List" and load a properly formatted inclusion list (see Figure 3 and section 4.1 for formatting on the inclusion list). The message "Inclusion loading was successful!" should appear in the console message box.

| | A | B | C | D | E | F | G | H | I | J |
|---|---|---|---|---|---|---|---|---|---|---|
| 1 | Parent Ion | Parent Charge | Retention Time Low | Retention Time High | Triggered Mass | Parent Ion Fragments | | | | |
| 2 | 577.8103 | 2 | 2.7 | 3.2 | 400.7014 | 239.1134 | 568.8055 | 333.2126 | 916.5084 | |
| 3 | 488.7267 | 2 | 2.7 | 3.2 | 616.8133 | 701.3192 | 630.2824 | 573.2615 | 458.2348 | 343.2081 |
| 4 | 381.8759 | 3 | 3.3 | 3.7 | 497.2526 | 902.472 | 773.4288 | 674.3607 | 551.3183 | 511.2979 |
| 5 | 403.2339 | 3 | 3.6 | 4.2 | 604.3472 | 908.5052 | 851.1824 | 752.4147 | 681.3772 | 496.2991 |
| 6 | 654.3091 | 3 | 4.1 | 4.5 | 981.4625 | 1159.5387 | 1088.5023 | 932.4495 | 888.3864 | 559.2279 |
| 7 | 513.3074 | 2 | 4.5 | 4.8 | 750.3401 | 912.5264 | 855.5032 | 685.3976 | 628.376 | 428.261 |
| 8 | 488.2779 | 2 | 4.8 | 5.2 | 356.5443 | 862.4655 | 765.413 | 652.3288 | 553.2976 | 431.7366 |
| 9 | 454.2646 | 2 | 5.3 | 5.8 | 670.6352 | 836.4861 | 765.449 | 666.3805 | 565.3337 | 365.2178 |
| 10 | 737.9198 | 2 | 5.3 | 5.8 | 1005.4503 | 1276.731 | 1163.6508 | 1064.583 | 879.5031 | 582.3345 |
| 11 | 713.364 | 2 | 7.5 | 7.7 | 543.3013 | 1149.5768 | 1062.5457 | 965.4925 | 850.4667 | 531.7762 |
| 12 | 895.9474 | 2 | 7.5 | 7.8 | 959.5194 | 1298.6666 | 1201.6143 | 1086.5873 | 901.5084 | 802.4395 |
| 13 | 1204.0403 | 2 | 7.8 | 8.2 | 1284.0854 | 1718.8663 | 893.4712 | 822.434 | 652.3276 | 537.3022 |
| 14 | 1004.4476 | 2 | 7.8 | 8.2 | 879.4848 | 1216.5499 | 386.1374 | 499.2212 | 702.3935 | 580.2673 |
| 15 | 919.3987 | 2 | 7.8 | 8.2 | 810.3566 | 1337.6163 | 1266.5803 | 1195.5436 | 983.3909 | 884.3224 |
| 16 | 803.0292 | 3 | 7.8 | 8.2 | 697.3751 | 1440.7379 | 1341.605 | 1142.6755 | 442.2762 | 626.3972 |
| 17 | 731.9089 | 2 | 7.8 | 8.2 | 637.8643 | 1150.5867 | 1053.5349 | 956.4828 | 784.3965 | 575.7966 |

*Figure 3. Example of a properly formatted inclusion list that is uploaded to the Moonshot Workflow by using the "Selection Inclusion List" button from Figure 1*

4.1. Inclusion List Columns and formatting: the headers "Parent Ion, Parent Charge, Retention Time Low, Retention Time High, Triggered Mass, Parent Ion Fragments" must be included. File should be saved as a tab-limited text file. Other delimiter, e.g. commas, are not supported. Each row represents a targeted ion.

- **1: Parent Ion.** The precursor mass list. These precursor values should be the mass of the internal standard peptides that have been added to the sample. If one of these parent ions is present within an MS1 scan (within the retention time window, see below), it will trigger a customized MS2 scan with isolation width, NCE, and first mass as defined by the MS2 properties. If more than one parent ion is present in an MS1, a multiplexed (MSx) MS2 scan will be sent with all present parent ions co-isolated and co-fragmented
- **2: Parent Charge.** The charge state of the parent ion. For a customized MS2 scan to successfully be triggered, the parent ion's charge state must match the charge state outlined in this column.
- **3: Retention Time Low.** The lower boundary of the retention time search for this specific parent ion. An ion present in an MS1 scan that has the parent ion mass (column 1) and parent charge (column 2), but that occurs before this retention time will not trigger a customized MS2 scan.
- **4: Retention Time High.** The upper boundary of the retention time search for this specific parent ion. An ion present in an MS1 scan that has the parent ion mass (column 1) and parent charge (column 2), but that occurs after this retention time will not trigger a customized MS2 scan.
- **5: Triggered Mass.** The triggered mass associated with the parent ion. Upon triggering a customized MS2 scan for the parent ion, this scan is analyzed for the known fragments (see Parent Ion Fragment Ions below). If the customized MS2 scan shows the required number of matched fragment ions, it will trigger another customized scan whereby the parent ion and the trigged mass are co-isolated and co-fragmented.
- **6: Parent Ion Fragment Ions.** The known fragments of the parent ion. These fragment masses will be utilized when the Moonshot Workflow analyzes customized MS2 scan of the parent ion/s. This field can have multiple columns associated with it where the number of known fragment ions is not limited. Each fragment ion should have its own column.

5. Press “Select MS2 Properties” (Figure 4). Set the “Defined First Mass”, “NCE”, and “Isolation Width”, “AGC Target”, “Max IT (ms)”, “MS Trigger Intensity Threshold” and “Dynamic Exclusion (s)” as per the requirements of your experiment. Press “Update MS2 Properties”.

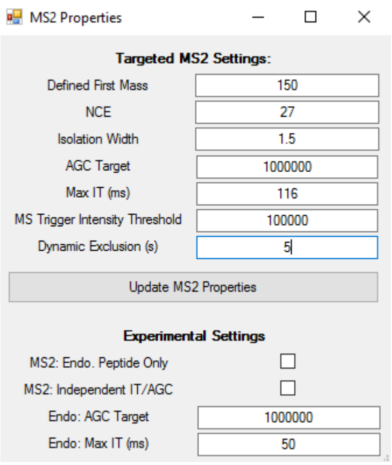


*Figure 4. MS2 settings screen that appears after clicking on Select MS2 Properties in the main screen (Figure 1)*

6. Modify the "PPM" and "Time (min)" in the main API screen (Figure 1) as per the requirements of your experiment, if needed. "Time (min)" should exceed the entire queue length, including loading steps. 'PPM' is the mass tolerance setting applied for the parent ions, triggered mass, and parent ion fragment ions in the inclusion list.
7. Start LC Queue, and immediately afterwards press Start on the API.
8. Once the acquisition of the raw file starts, the API will start to monitor the targets (parent ions) in the inclusion list. When a precursor mz in the inclusion list is found in the corresponding time frame and above the MS intensity threshold, the console output will the updated. Figure 5 shows an example of console output.

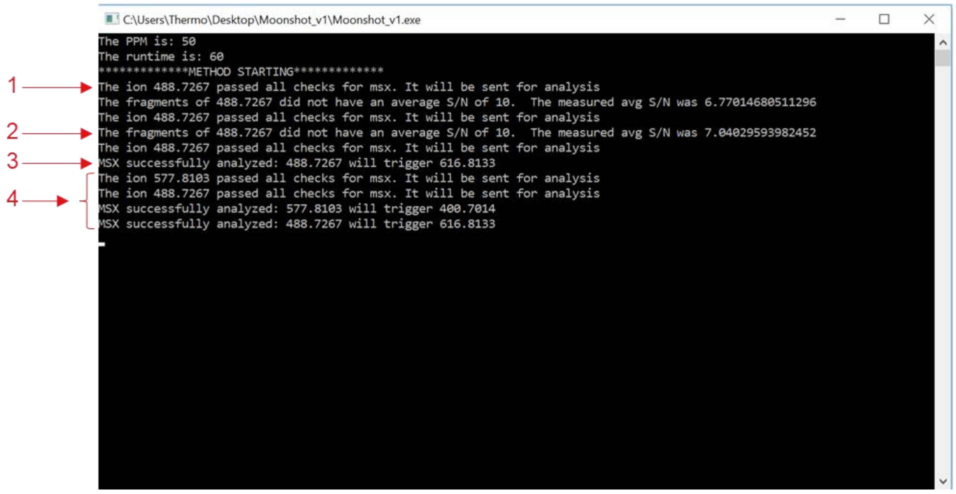


*Figure 5. Secondary console window showing messages during MS acquisition in a hybrid-DIA run.*

**#1:** The message indicates that the parent ion 488.7267 was present with the retention time window defined by the lower and upper retention boundary conditions in the inclusion table, i.e. Retention Time Low and Retention Time High, respectively. This mass will then be isolated with a narrow isolation window (as defined by the Isolation Width of the MS2 Properties) and fragmented in a customized MS2 scan.

*Scans:* *Full MS1 → MS2 of 488.7267*

**#2:** The message indicates that the customized scan of 488.7267, which was generated by Example 1, showed some known fragment ions (see “Parent Ion Fragment Ions” in section Format of Inclusion List); however, the average signal-to-noise (S/N) was not high enough (>10) to trigger an MSX MS2 with the parent and triggered masses.

*Scans: Full MS1 → MS2 of 488.7267 → Analysis failed so: Continuation of normal method scans (e.g. DIA Windows, Full Scan etc.)*

**#3:** The message indicates that a customized scan of 488.7267 (a different customized scan than generated in Example 1 and analyzed in Example 2) was successfully analyzed, meaning the known fragment ions were present at a S/N of >10, therefore a MSX where the parent ion, 488.7267, and the triggered mass (see “Triggered Mass” in section Format of Inclusion List) 616.8133 will be co-isolated and co-fragmented

*Scans: Full MS1 → MS2 of 488.7267 → Analysis passed so: MS2 MSX of 488.7267 and 616.8133 → Continuation of normal method scans (e.g. DIA Windows, Full Scan etc.)*

**#4:** This message block shows the entire workflow message output when more than one Parent Ion is present within an MS1 scan. Both 577.8103 and 488.7267 were present in the MS1. An MSX where both of these masses were co-isolated and co-fragmented was generated. Analysis of this MSX scan showed the fragments of both parent ion present; therefore, two customized scans are sent. The first is an MSX scan that co-isolated 577.8103 and its corresponding triggered mass of 400.7014. The second is an MSX scan that co-isolated 488.7267 and its corresponding triggered mass of 616.8133.

Scans: Full MS1 → MS2 MSX of 577.8103 and 488.7267 → Analysis passed so: MS2 MSX of 577.8103 and 400.7014 → Analysis passed so: MS2 MSX of 488.7267 and 616.8133 → Continuation of normal method scans (e.g. DIA Windows, Full Scan etc.)

9. Two files will be generated during hybrid-DIA acquisition and they are located in “C:\ProgramData\Thermo\Exploris\Instrument\Moonshot”
   9.1. A log-file containing the data from the API acquisition (e.g. 2022_12_14_10_15 Moonshot Workflow.txt).
   9.2. An accessory file with all information from each full scan (Scan number, mz, charge, intensity and resolution). This file can be used for troubleshooting purposes. It is named with a time-stamp followed by the name of the corresponding raw file. These files can be deleted if the space in the disk is limited.

**Supplementary Data 2.** Guide for multiplex IS/ENDO extraction and data analysis is hybrid-DIA files.

### Extraction of MSx scans: MSx_Extractor.py[1]

Software required:

Python version: 3.9.7 (https://www.python.org/downloads/release/python-397/), Proteowizard version 3.0.21246 (https://proteowizard.sourceforge.io/)

Python Packages required: tkinter, sys, time, os, pymsfilereader, pathlib[1]

**INPUT**:

- Dir: folder with raw files to convert.
- IS-ENDO file: pairs of mz for IS and ENDO peptides (Figure 1)

```
434.881286,432.209886
1009.486558,1004.482423
1044.976839,1040.969739
1048.145059,1045.473659
1189.560505,1186.224415
1206.039222,1201.035088
367.931562,365.429495
408.211336,404.204237
409.203661,405.867572
424.722068,420.714969
448.194502,444.187402
```

*Figure 1. Example of IS-ENDO File*

**OUTPUT**:

- Mzml files: one mzml file per raw file containing only the desired MSx scans. To be used as input for Skyline.
- Results.txt files: one per raw file, contains the injection times per scan. Required for IT normalization. Copy all txt files into a new folder and label it as “txt”.
- Config.txt: one file per raw file, used internally by ProteoWizard. They can be deleted when the conversion is finished.

**BEFORE START:**

1. Locate the folder that contains the installation of ProteoWizard.
2. Open with a text editor (such as NotePad++) the app “MSx_Extractor.py”
3. Go to line 62 in the python script and update the directory to the one where you have installed ProteoWizard. Remember to keep the “\\” between subdirectories.
4. Copy all the raw files into a new folder.

**RUNNING THE APP:**

1. Double click on MSx_Extrator.py. Two windows would appear (Figure 2 and 3), one corresponding to the GUI and the python console.
2. Upload IS ENDO file by clicking on “Open file” and navigating to the location of the file.
3. Indicate the folder that contain the .raw files of interesting by clicking on “Select Dir”.
4. Click “Go!” to start the analysis. A message saying “Analyzing sample n/n” would appear and it would be updated every time a raw file is done. When all raw files have been processed, the message “Done” will appear.
5. During the raw file processing the python console (Figure 3) would show ProteoWizard actions.

[1] For help installing python packages: https://packaging.python.org/en/latest/tutorials/installing-packages/#id18

6. When done, three files would have been generated per raw file: .mzml, results.txt and config.txt. For convenience in further steps, copy all results.txt files into a new folder.

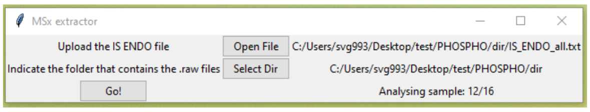


*Figure 2. MSx_Extractor.py main window.*

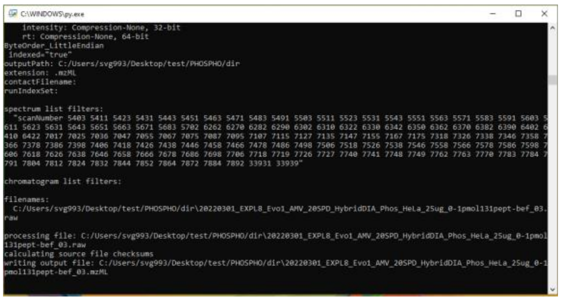


*Figure 3. Python console.*

### Import into Skyline: Extraction of the transitions.

Required: Skyline template with the desired transitions (.sky) and fragment library (.blib).

1. Edit the following parameters in the "Transition settings tab" (see figure below for details):
    - Filter tab: remove "b" ions from ion types. They cannot be used for quantification since they are shared between the heavy and light peptides.
    - Instrument tab: set "method match tolerance m/z" to 0.001.
    - Full-Scan:
        - MS1 filtering: none
        - MS2 filtering: PRM

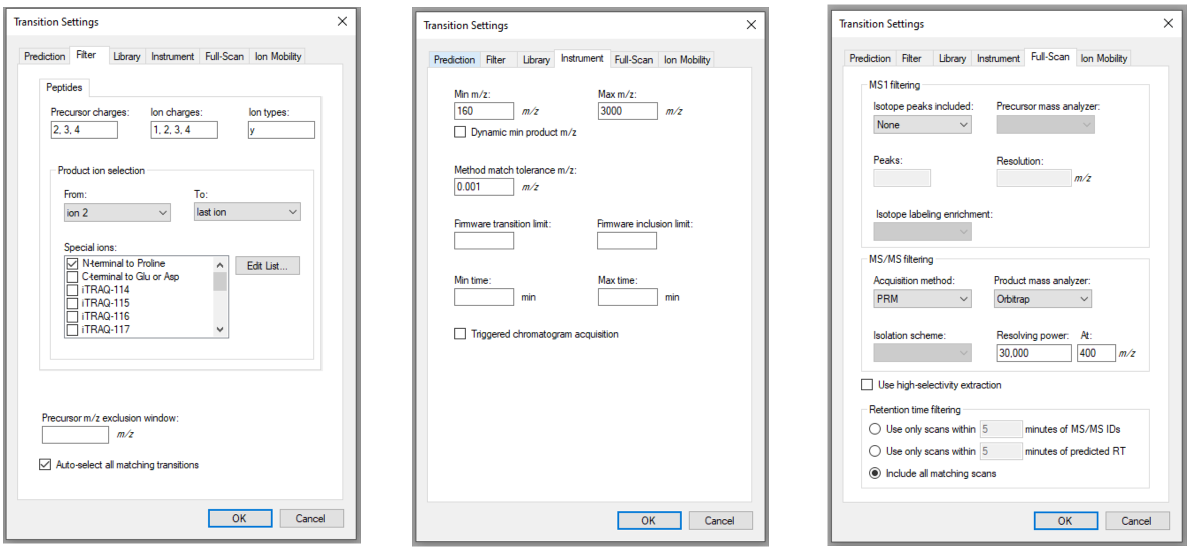


*Figure 4. Transition settings tab in Skyline with specific parameters required for the succesfull import of the processed mzml files.*

2. Import mzml files: File > Import > Results
3. Open Document grid (Alt+3 or View>Document Grid). Edit a document to contain the following information and save the report as "Quantification_IS_ENDO_forHybrid".
    a. Peptide.
    b. Protein. IMPORTANT: This header would indicate in the next step the IDs for each IS/ENDO pair. In the case of more than one peptide per protein, modify the header to "ProteinName", and rename "Peptide" column to "Protein".
    c. Replicate.
    d. Raw Intensities.
    e. Raw Times.
    f. Fragment Ion.
    g. Product Mz.
    h. Isotope Label Type.
    i. Mass Error PPM.
    j. Peptide Peak Found Ratio.
    k. Raw Spectrum Ids.
4. Export Report "Quantification_IS_ENDO_forHybrid" (.csv format).

5. Open Document grid again and load the table named “All_Precursors_Initial_Survey”. Edit the table to contain only two columns:
    a. Protein Name. IMPORTANT: This header would indicate in the next step the IDs for each IS/ENDO pair. In the case of more than one peptide per protein, choose the appropriate column for each unique pair (i.e: Modified Peptide Sequence). The content needs to match the content of the “Protein” column from the Quantification report.
    b. Precursor Mz.
6. Export Report “All_Precursors_Initial_Survey” (.csv format).

### Injection time normalization, quantification and visualization of results.

Software required: R (v4.0)
R packages required: shiny, shinyFiles, shinycssloaders, dplyr, ggplot2, data.table, gridExtra, tidyr, ggpubr, MESS, config

1. Copy the file app.R and config.yml to your computer into a folder called “Shiny-APP”. The file “config.yml” contains certain parameters to filter the data do the quantification. There data in the file can be used as a default, but it can be modified if needed. The parameters are:

   ppm: max error mass allowed. Default: 10 ppm
   n_y: min number of y fragments required for quantification. Default: 1
   ppfr: peptide peak found ratio from Skyline. Default: 0.5

2. Open R (it can be the R console or Rstudio) and type:

```
library(shiny)
setwd("C:/Users/xxxx/xxxx/dir_shiny_app/")
#Indicate the location of the folder "Shini-APP"
runApp("Shiny-APP")
```

   or click on “Run App” on the right top corner.

3. A shinyApp will open in a new window (Figure 5 and 6).
   Input:

   (1) directory where the txt files generated in the first step are stored.
   (2) common pattern in the raw files. If the raw files share a common pattern and it has been removed during Skyline processing, indicate it here.
   (3) Maximum injection time (IT) used during hybridDIA acquisition
   (4) Precursor MZ list: csv file that contains the mz for the heavy and light peptides together with their id (phospho-site, peptide).
   (5) Quantification results: csv files exported from Skyline in the previous step. When this document is loaded, two selection menus will appear, one for the sample and another for the peptide/phospho-site.

   Output:

   (6) Plot phospho-site: this button will plot the XIC (6a) of the IS and ENDO peptide for the selected phospho-site in the selected sample before and after IT normalization. Also, it will show a profile plot (6b) of the relative intensity (calculate from the ratio IS/ENDO) of that peptide across samples.
   (7) Print Heatmap: this button will plot the relative intensity (calculates from the ratio IS/ENDO) of all peptide across samples in the form of a heatmap. (In the “All results” tab).
   (8) AUC IS, AUC ENDO: these buttons will export a table with the AUC for the heavy (IS) (8b) and light (ENDO) (8a) counterparts for all peptides, normalized by IT.

(9) If this check-box is active, the profile plot will show the ratio IS/ENDO (heavy to light) and not the scaled values across replicates. Also, the heatmap will be plotted using the ratio IS/ENDO.

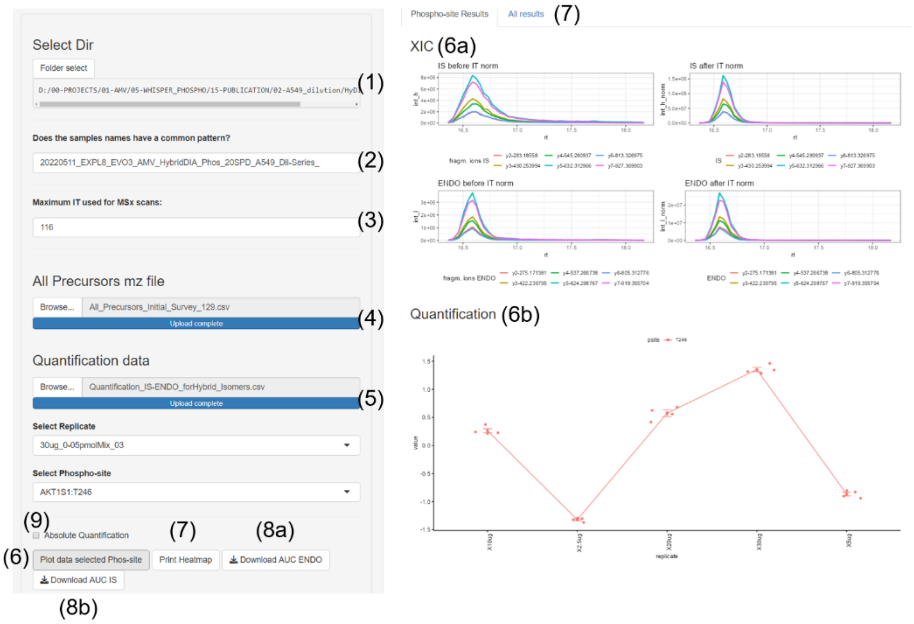


*Figure 5. Shiny App interface. On the left panel: option menu to upload required files. Once Quantification data file is uploaded the selection tabs "Selection Replicate" and "Select phospho-site" appears for the user to choose what to plot. On the right panel: (top) XIC of peptide AKT1S1-T246 for sample "30ug, replicate 3", (bottom) relative quantification of that peptide among the samples compared.*

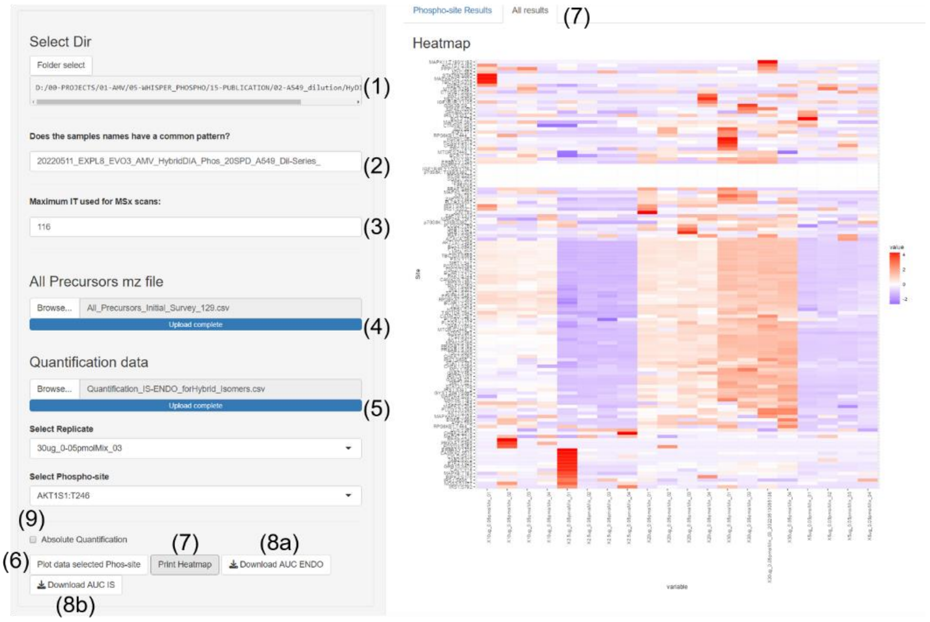


*Figure 6. Shiny App Interface. On the right panel: Heatmap with the relative quantification across samples of all peptides monitored in the loaded hybrid-DIA experiment.*

Suppl. Figure 1 Martinez-Val et al.

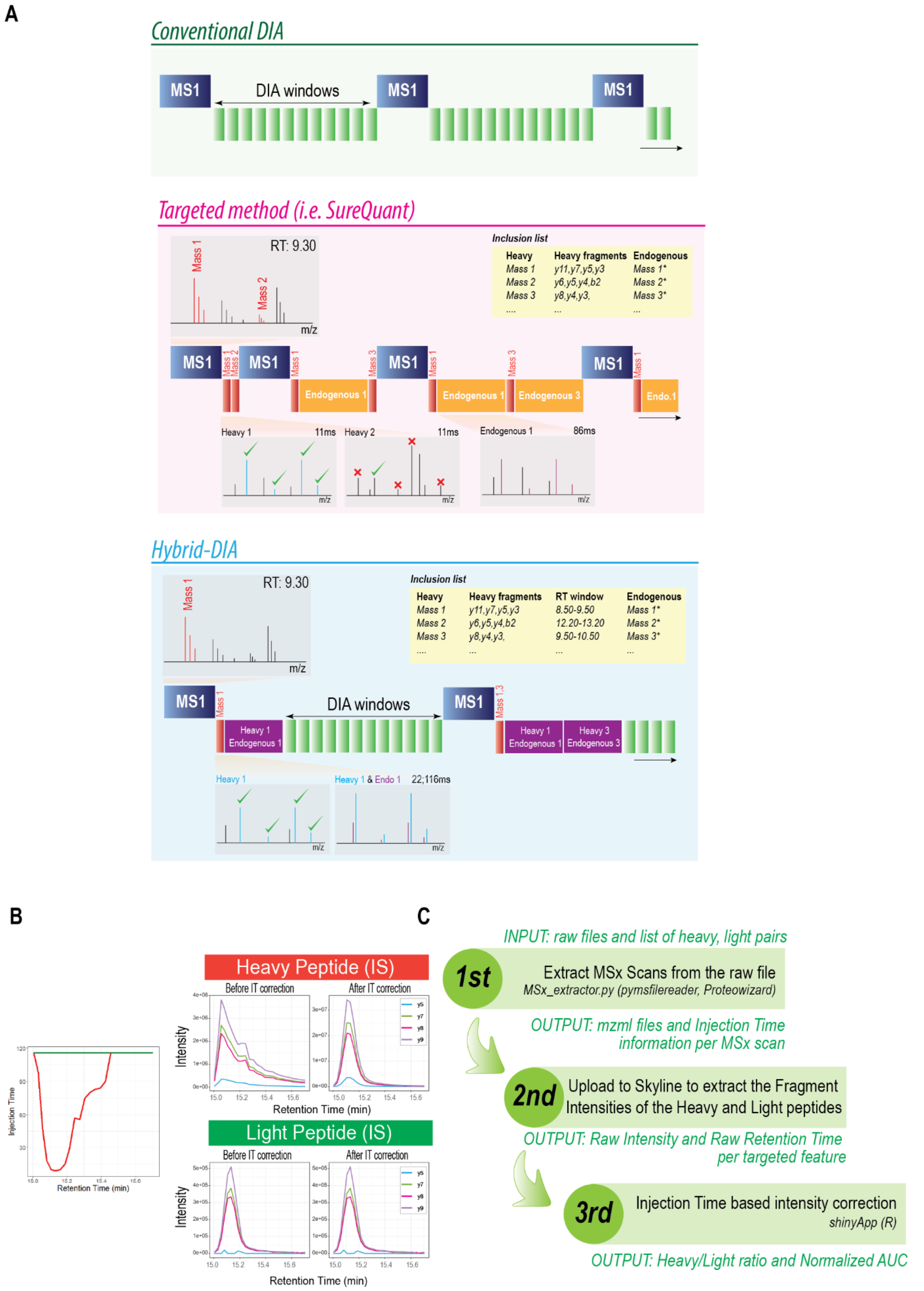


**Supplementary Figure 1.** (A) Schematic diagram of the different acquisition methods used in this project: DIA, spiked-in triggered targeted methods and hybrid-DIA. (B) Example of the effect of the MS2 intensity profile due to the differential injection time for heavy and light used in multiplexed scans and the result after correcting the MS2 intensities by that injection time. (C) Diagram of the three steps of the data analysis pipeline for extracting and processing the targeted multiplexed scans generated with the hybrid-DIA method.

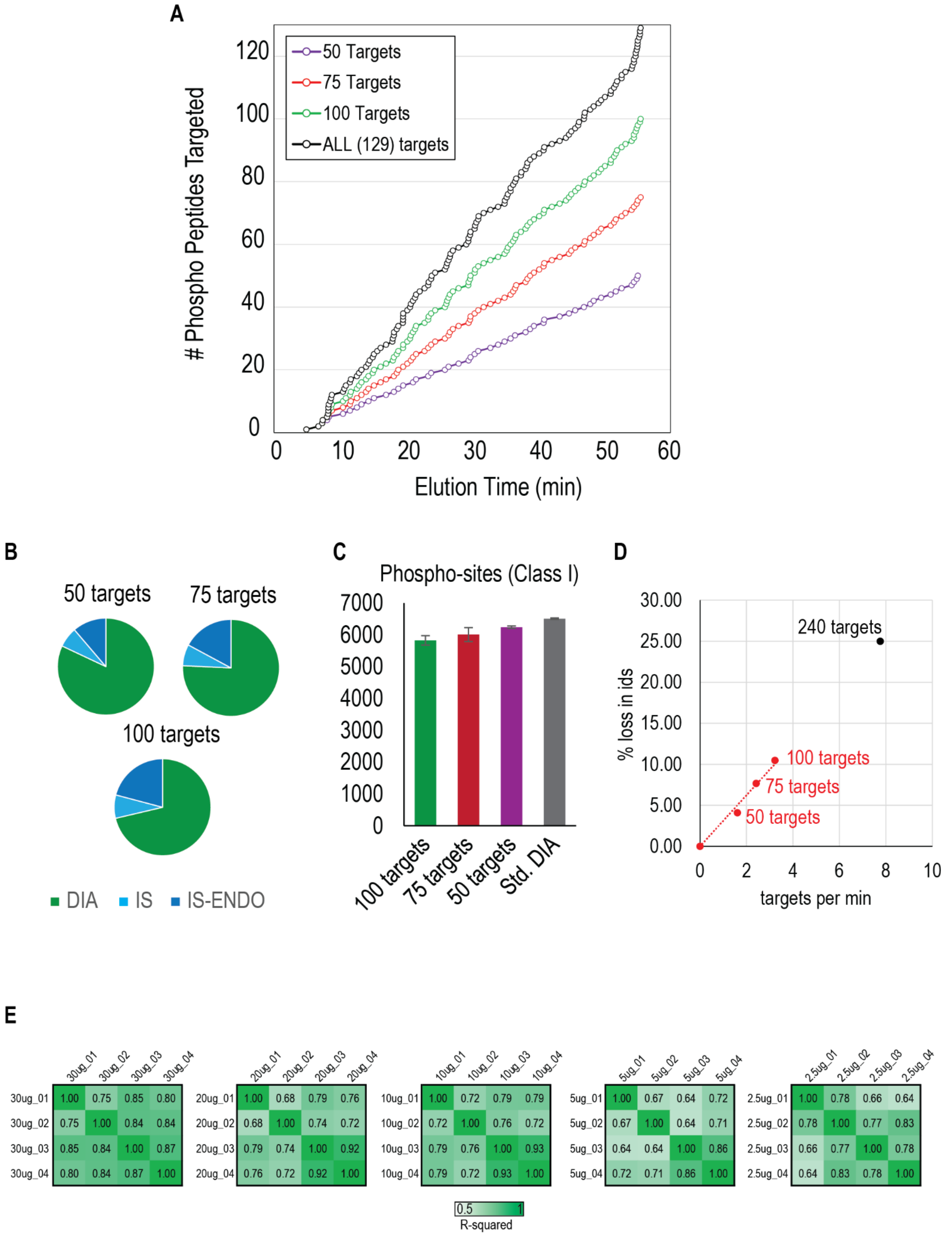


**Supplementary Figure 2.** (A) Graph representing the distribution of the phospho-peptides from the SureQuant Multipathway Phosphorylation Kit (black), and the different subset of targets used for evaluating cycle time use: 50 (purple), 75 (red) and 100 (green). (B) Pie charts representing the proportion of cycle time used by the hybrid-DIA API measured by acquisition time when using an inclusion list of 50, 75 or 100 targets. In green, the proportion of the total MS2 acquisition time used in DIA scans; in light blue, the time used in survey scans for detecting the presence of the internal standard (IS) and in dark blue, the time used in multiplexed (IS and ENDO peptides). (C) Bar plots showing the number of phospho-sites (class I) identified in either standard DIA runs, or when using hybrid-DIA methods and increasing number of targets. Bar length represents the average of the identified sites between replicates, and error bars the standard deviation (n=2). (D) Relationship between peptides targeted per minute (calculated by dividing the total number of peptides in the inclusion list by the gradient length) against the percentage of lost identifications (using standard DIA values as reference). In black, hypothetical extrapolation of the number of targets that would lead to 25% of lost identifications. (E) Sample to sample correlation between DIA runs, measured as R-squared.

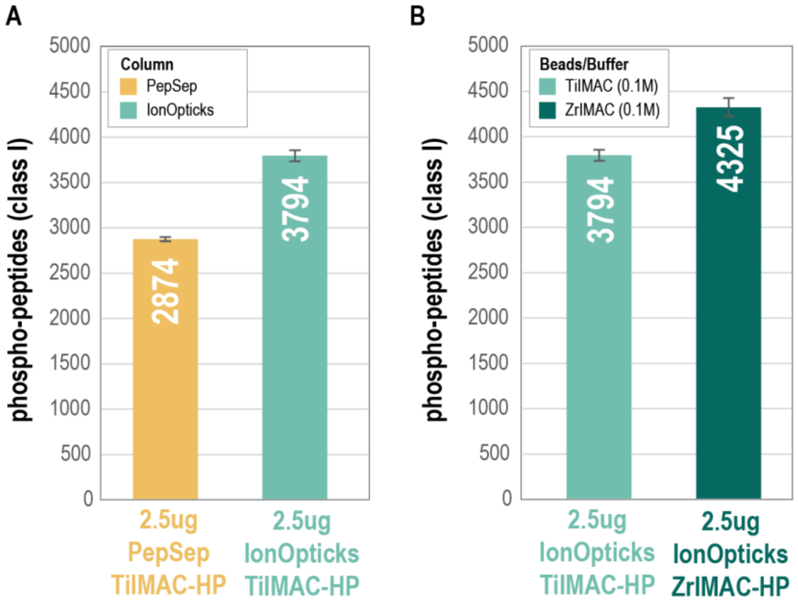


**Supplementary Figure 3.** (A) Comparative bar plot assessing the effect of the column type (PepSep vs IonOpticks) on the number of phospho-peptides (class I) identified when using 2.5ug of peptide input for phospho-enrichment with TiIMAC-HP beads (n=3). (B) Comparative bar plot assessing the effect of the beads type (TiIMAC-HP vs ZrIMAC-HP) on the number of phospho-peptides (class I) identified when using 2.5ug of peptide input for phospho-enrichment with IonOpticks column (n=3).

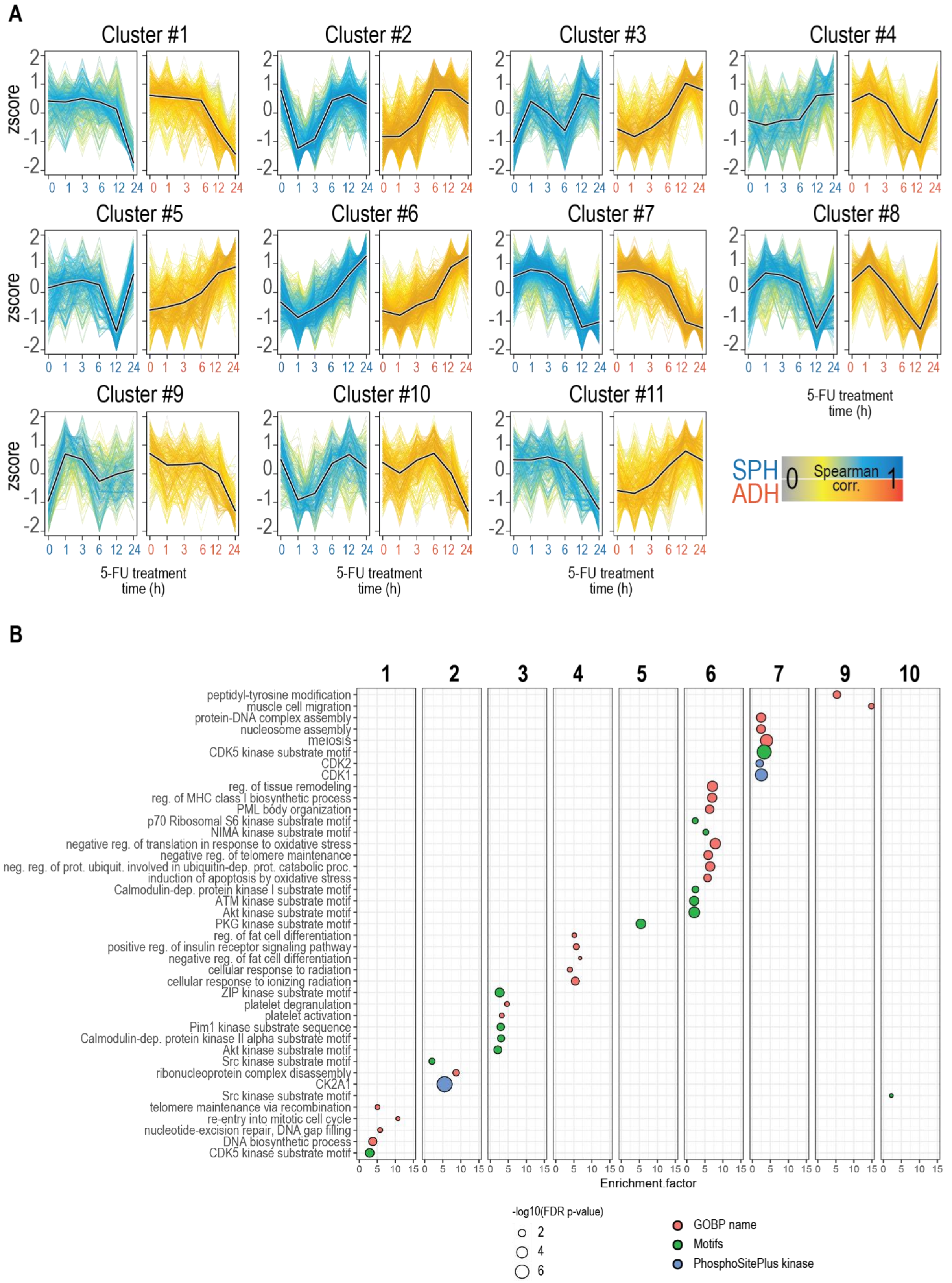


**Supplementary Figure 4.** (A) Temporal profile clusters for Spheroids (blue) and Adherent (yellow) cells upon 5-fluorouracil treatment. Y-axis represents z-score scaled log2 phospho-site intensity. Dark line indicates the centroid of the cluster, and each line represents a different phospho-site. Color of the lines reflects the spearman correlation to the centroid of the cluster.

(B) Fisher's exact test to evaluate enrichment of GO-BP terms, kinase motifs and PhosphoSitePlus kinases in the clusters defined in panel A. Only clusters with relevant GO-BP terms are used in this panel.
